# Generative and integrative modeling for transcriptomics with formalin fixed paraffin embedded material

**DOI:** 10.1101/2025.02.21.639356

**Authors:** EJ Mucaki, WH Zhang, A Saha, S Trabinjac, S Nofech-Moses, E Rakovitch, V Dumeaux, MT Hallett

## Abstract

Formalin-fixed paraffin embedded (FFPE) samples are challenging to profile using existing high-throughput sequencing technologies, including RNA-seq. This difficulty primarily arises from the degradation of nucleic acids, a problem that becomes particularly acute with samples stored for extended periods. FFPE-derived RNA-seq (fRNA-seq) data have a high rate of transcript dropout, a property shared with single cell RNA-seq. Transcript counts also have high variance and are prone to extreme values. We introduce the PaRaffin Embedded Formalin-FixEd Cleaning Tool (PREFFECT), a probabilistic framework for the analysis of fRNA-seq data. PREFFECT uses generative models to fit distributions to observed expression counts while adjusting for technical and biological variables. The framework can exploit multiple expression profiles generated from matched tissues for a single sample (e.g., a tumor and morphologically normal tissue) in order to stabilize profiles and impute missing counts. PREFFECT can also leverage sample-sample adjacency networks that assist graph attention mechanisms to identify the most informative correlations in the data. We demonstrate how PREFFECT uses this information to more accurately compute downstream analyses such as sample clustering in several datasets.

## Introduction

Formalin-Fixed Paraffin Embedded (FFPE) material has long been used in histopathology to store samples in a manner which preserves cellular structure and tissue morphology. The use of FFPE blocks for immunohistochemistry has been used in diagnostic pathology since 1991 [1]. Moreover, FFPE material is a significant resource for retrospective clinical research and provides utility in prospective studies, potentially eliminating the need for the collection of fresh frozen specimens. The development of antigen retrieval techniques for extracting nucleic acids and proteins has broadened its range of applications to include, for example, Western blot analyses, mass spectrometry, and PCR. Its ease of storage and cost-effectiveness has enabled the cataloging of the pathology of health and disease [2] and several hundred million archival FFPE samples exist worldwide [3].

FFPE-harvested material can be of sufficient size and quality for use in modern -omic projects [4–8] including RNA-seq [9–16], which is the primary focus of our effort here. The use of these technologies, however, presents significant challenges due to technical artifacts that arise due to the preparation and storage of the material. For example, when the tissue is fixed with formalin, methylene bridges can form between amino groups, altering the structure of nucleic acids and proteins by adding reactive groups [17], resulting in fragmentation and mutations [18]. The dehydration process can cause denaturation of nucleic acids and proteins, including the loss of disulphide and hydrophobic (for proteins) bridges, and hydrogen bonds that maintain secondary and tertiary structure. RNA stability may be greatly reduced after renaturation. Heat, modulation of pH levels and endogenous enzymes (e.g., nucleases) during de-crosslinking can cause further nucleic acid damage particularly to labile RNA [19]. The rate of degradation of nucleic acids is dependent on time, the fixation process used, and storage conditions [20–22]. Extraction, which requires methods to generate thin sections of tissue for staining or tissue ribbons for nucleic acid extraction, also causes damage through physical shearing of the nucleic acids and oxidative damage due to the use of, for example, phenols. Hydrolytic deamination can occur during several of the steps described above and can result in the conversion of cytosine into uracil [15, 23, 24] or guanine to adenine [25]. NGS-based studies of FFPE material consistently identify a significantly higher mutation rate compared to fresh frozen tissue [25–27] and measures of RNA quality (e.g. RNA integrity number, DV200) are significantly lower for FFPE-derived material compared to fresh frozen tissue [28].

The quality of FFPE-harvested material has improved due to advancements in protocols for nucleic acid extraction, sequencing preparation, and bioinformatic analyses [4, 16, 29–31]. Comparisons between fresh frozen and FFPE-derived tissue regardless of extraction and sequencing preparation methods have shown that many transcripts have zero counts in the FFPE RNA-seq (fRNA-seq) profiles; these zero values are likely due to technical problems and do not represent a true biological expression level [4, 8, 27]. Such observations underscore the need for specialized normalization/de-noising methods for fRNA-seq profiles such as those proposed by Yin and colleagues [32, 33] where count data is fit to a two-component model comprising a truncated normal distribution (for expressed transcripts) and a zero-inflated Poisson distribution (for non-expressed transcripts). Efforts to precisely characterize the distributional properties of fRNA-seq profiles nevertheless lag significantly behind other transcriptome profiling areas, including single cell RNA-seq (scRNA-seq) where high dropout rates are also observed.

This effort starts with investigating the distributional qualities of the fRNA-seq data. In particular, we examine some of the underlying technical and biological variables affecting transcript count data and provide evidence that it is well-modeled by the negative binomial (NB) distribution, a property shared with bulk and scRNA-seq data. There are, however, challenges unique to fRNA-seq data including observations that transcript counts are sometimes four orders of magnitude greater than scRNA-seq and with very high variability and dispersion. Moreover, the relatively small sample size in FFPE studies makes fRNA-seq analysis potentially an even greater challenge than scRNA-seq: the estimated 1500 fRNA-seq datasets in international repositories (e.g., ENA, SRA, dbGaP) typically contain on the order of 10^2^ samples, while most scRNA-seq studies contain on the order of 10^4^ − 10^6^ samples. In response to these challenges, we introduce the PaRaffin Embedded Formalin-FixEd Cleaning Tool (PREFFECT) to provide a probabilistic model for fRNA-seq data. PREFFECT uses generative models to re-express observed transcript counts using either an NB distribution or a zero-inflated extension alongside metadata corresponding to known technical and biological effects in the data. Similar approaches have been explored in the context of scRNA-seq data analysis [16, 34–51]. PREFFECT contains a series of conditional variational autoencoders (cVAEs) which allow multiple tissues to be considered simultaneously in addition to graph attention mechanisms [47, 52] which assist the learner to borrow information across samples and tissues. The goal is to build latent spaces that factorize expression profiles into their biological and technical variables, allowing for the ablation of nuisance parameters and tempering of extreme measurements, thereby providing a more refined means to perform common downstream analyses including clustering, differential expression, and survival analyses. We explore the use of PREFFECT with several publicly available datasets.

## Results

### FFPE-derived RNA-seq expression is well-modeled by the negative binomial distribution

To characterize the distribution of transcript counts obtained from whole transcriptome fRNA-seq profiling, a compendium was constructed by selecting publicly available datasets which contain a considerable number of profiles (*≥*20) and for which the corresponding raw, non-normalized count data is available. The *N* = 13 sets of fRNA-seq samples vary in terms of tissue types, cell types, disease (or normal samples), age of samples, and technical variables (Supplementary Table S1). The transcripts with the highest counts tend to be the same as those observed in scRNA-seq studies including *MALAT1, NEAT1, XIST* [53] *and others listed in Figure 1A. These extreme counts in the right tail induce a very high standard deviation with respect to the average count per transcript (Figure 1B). For example, the mean count for MALAT1* in the TMBC dataset [54] is *∼* 4.4 million but the maximum observed value is 34 million. Mean trimming of the right-most 1% of extreme observations reduce the overall standard deviation by an order of magnitude in almost all datasets (Figure 1A). However, it should be stressed that even after trimming, the range of counts for many transcripts in fRNA-seq data remains very broad. For example, transcript counts for the *estrogen receptor 1 (ESR1)* in the Sunnybrook fRNA-seq dataset range from just 1 to more than 100, 000.

**Figure 1.**
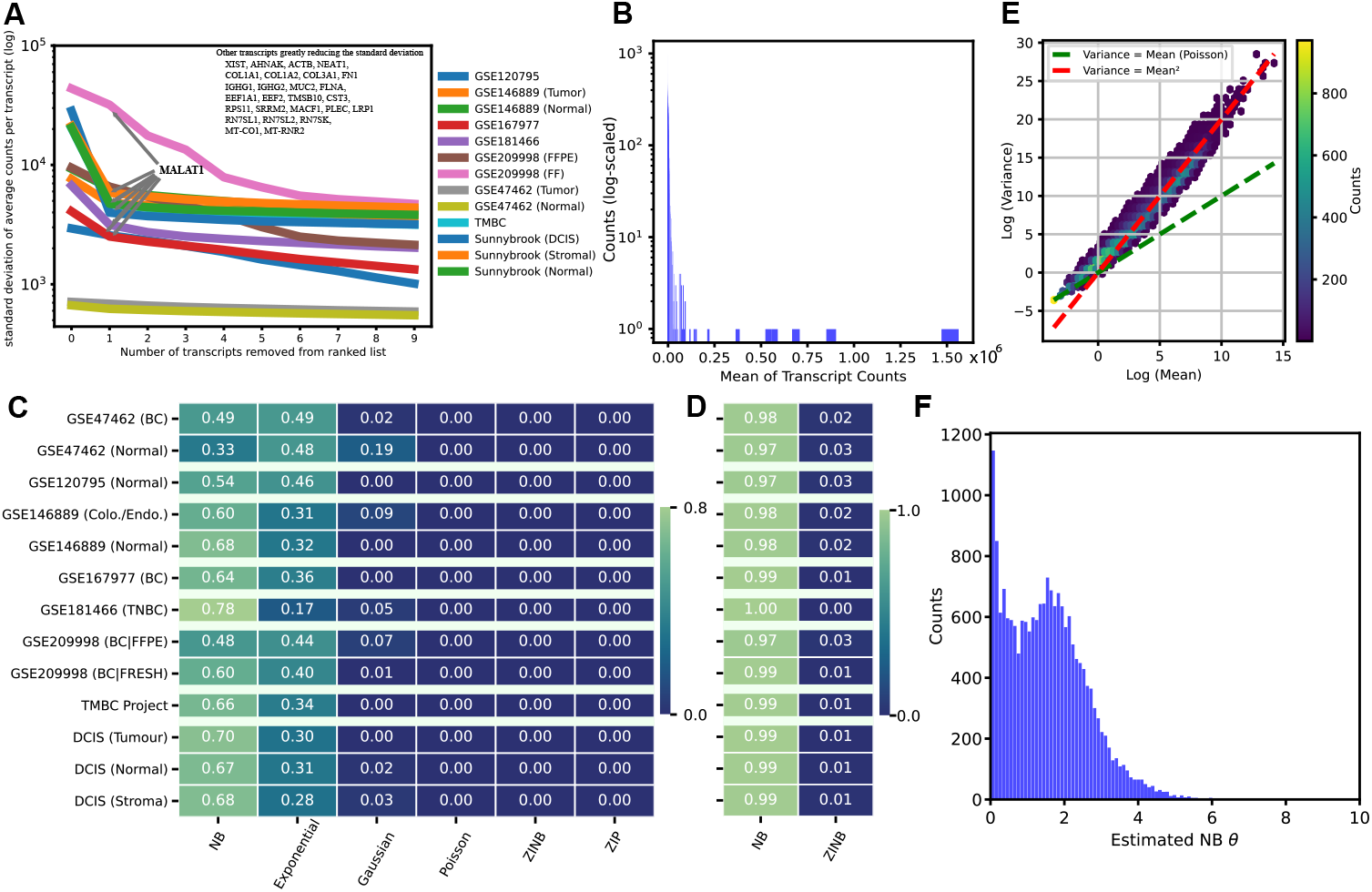
Distributional properties of fRNA-seq data. **A**. For each dataset in our compendium, transcripts were ordered from highest to lowest count. Depicted here is the number of highest transcripts removed from each dataset versus the reduction in the standard deviation. *MALAT1, NEAT1, XIST, COL1A1/2, MT-* mitochondrial and *RN-* ribosomal genes are highly and ubiquitously have many reads likely due to their large size. The standard deviation drops by an order of magnitude after the highest 5 transcripts are removed. **B**. Histogram of the (log) mean count for transcripts in the *GSE209998* dataset. Note the extreme outliers *>* 10^5^ counts. **C**. For each dataset (row), the fraction of all transcripts with the best fit by distribution (column). The NB distribution has the highest frequency for all but *GSE47462*. **E**. The vast majority of transcripts are better fit by the NB than the ZINB under the AIC criterion. **E**. Hexagonal heatmap of the log mean versus the log variance of each transcripts in *GSE209998*. The green line indicates the trend if variance equaled the mean, while the red line indicates the trend for a quadratic mean-variance relationship. **F**. Distribution for estimated dispersion *θ* with dataset *GSE47462*.

Six different distributions were fit to the data (Methods 2.2) using the Akaike information criterion (AIC) as a measure of fit. A significant majority of transcripts are best fit by the negative binomial (NB) distribution for all but the normal tissue samples of GSE47462 (Figure 1D). Because the various distributions differ in their number of parameters and the AIC contains a weak penalty for model complexity, we repeated the analysis using the Kolmogorov-Smirnov *D* statistic, which does not adjust for model size, but arrived at the same result (Supplementary Information 3).

Figure 1D also suggests that transcripts often prefer the exponential distribution. However, we observed that this preference is highly dependent on the percentage used to trim outliers. That is, the preference for the exponential fit is strongly influenced by the number of highly expressed transcripts in the right tail (Supplementary Figure S1B), further highlighting the fact that fRNA-seq data is prone to extreme measurements.

A significant fraction of transcripts have zero counts in all of the fRNA-seq datasets (ranges from 0.1 to 0.5 with median 0.46; Supplementary Table S1). Zero-inflated (ZI) extensions to distributions like the NB are often considered in such cases. In addition to the mean *µ* and dispersion *θ* of the NB distribution, the ZINB has a third parameter *π* that controls the probability of a so-called *dropout event* (a zero count for a transcript that is not a variate from the NB). However, Figure 1D reports very little support for the ZINB. A direct comparison between the NB and ZINB reaffirms that observed 0 counts are almost always well-modeled by the NB alone (Figure 1E). Additional analysis in the Supplementary Information provides evidence that this is not due to model complexity. It is common in the context of NB-related distributions to express the variance as a function of the mean and the dispersion parameter *θ* as follows:

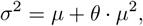

highlighting the fact that high values for *θ* induce a large variance in the NB distribution. This variance often appears to be sufficient to model the observed quantity of 0 counts in fRNA-seq data, a conclusion which has also been reached for scRNA-seq data [41]. Figure 1E reaffirms the choice of an NB over a Poisson distribution, since the variance is observed to be greater than the mean.

After fitting an *NB*(*µ, θ*) to each transcript within each dataset, the overall mean *µ* is observed to be 710 but with an extremely large standard deviation, ranging from 666 to 43, 713 (Supplementary Table S1). We also observe a mean dispersion *θ* of 1.47 with a large relative standard deviation of approximately 1.5. Few transcripts have an estimated *θ* below 0.01 and few transcripts have an estimated *θ* above 5. An NB distribution with *θ <* 1 is maximized at 0 with many near-zero counts, whereas larger *θ* are maximized strictly above 0 (see Figure 4C). Many transcripts, including *ERBB2/HER2* and *ESR1*, have larger values for *θ* greater than 3 (Figure 1F); these induce a flat, long NB distribution.

### fRNA-seq data as a mixture of distinct technical and biological effects

It is well-established that technical variables often have a significant impact on distributional parameters in -omic profiling. This includes variables such as library size, batch number and library complexity (e.g., percent duplication and DV200). Across the fRNA-seq compendium, library size was highly dependent on batch number in all datasets where this information is available (one-way ANOVA, all *p <* 0.001). Moreover, both the location *µ* and scale *θ* parameters for the fitted NB distributions are almost always significantly different between batches (log-likelihood-based test of the change in fit between batch-dependent parameters). The percentage of duplicate reads also differed significantly between batches (*p <* 0.01 for all available datasets).

Biological variables (e.g., hormone receptor status, proliferative index, grade in cancer studies) should of course have a significant impact on the fRNA-seq profiles. To investigate how count distribution is affected, we focused on patient subtype across the breast cancer datasets within the compendium. It is well established that breast cancer samples can be partitioned into at least five distinct subtypes at the transcriptional level. PAM50 is a commonly used classification tool that subtypes samples according to the counts of 50 specific transcripts for invasive [55] and in situ [56] lesions. Large differences in fitted NB parameters were observed when samples were stratified by the PAM50 subtype for the vast majority of the transcripts (Figure 2 and also Supplementary Figure S2). For example, the location parameter of the NB distribution for estrogen receptor 1 (*ESR1*) is markedly higher for luminal A (dark blue) and B (light blue) estrogen receptor positive subtypes in comparison to all remaining estrogen receptor negative subtypes; this is consistent with the original data represented Figure 2A. The fitted subtype NB distributions for Keratin 5 (*KRT5*) have the largest location parameter for normal-like lesions (green), the subtype where it is highest expressed according to the original PAM50 manuscript [55]. The NB fits for *ERBB2 (HER2)* and *GRB7*, two genes in the 17q12 amplicon characteristic of HER2 positive tumors (pink), have location (*µ*) and scale parameters (*θ*) well above zero only in the Her2 subtype. Note also that *ERBB2 (HER2)* counts are at least one order of magnitude larger than all other transcripts, and there is a large variance in counts across the samples. Likewise, epidermal growth factor (*EGFR*), a well established marker of basal-like (red) and normal-like (green) tumors have similar NB fits in these two subtypes. The NB distribution here is flat, which corresponds to a high *θ* as discussed in earlier.

**Figure 2.**
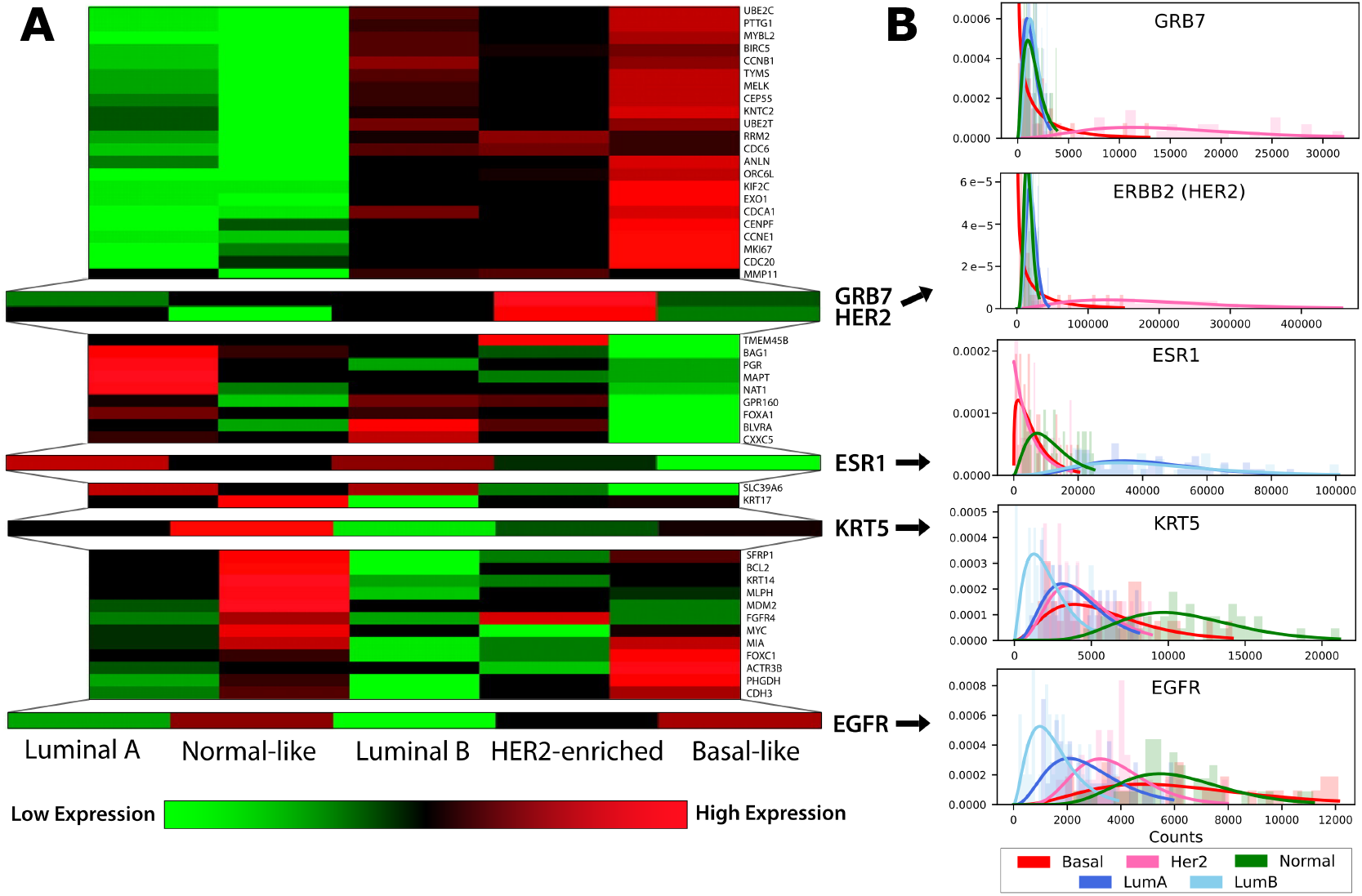
**A**. Heatmap modified from [55], depicting the expression of each of the PAM50 transcripts across the five breast cancer subtypes in the original data. Here, red and green depict over- and under-expression of the transcript respectively. **B**. Histograms of transcript counts in the Sunnybrook DCIS tumor cohort for selected PAM50 genes (enlarged rows of the heatmap in **A**) colored by their subtype. A NB distribution was fit for each transcript in each subtype independently after median library size adjustment and trimming.

We again do not see strong evidence for use of the ZINB over the NB distribution with any of the PAM50 genes when samples are stratified by subtype (Supplementary Figure S2). The need for the ZINB distribution would be clearly justified if we observed a large spike of zeros in cases where *µ* is well above 0. Instead, it appears that the NB distribution with a suitably high dispersion *θ* is sufficient in cases where *µ* is close to 0. We conclude that fRNA-seq datasets are well-modeled by the NB distribution albeit with large dispersion at times.

### The core generative model

PREFFECT is a collection of cVAEs designed to model transcript count data by fitting it to NB or ZINB distributions. Count matrices are expressed as a mixture of different technical and biological effects, allowing for the adjustment of data to account for factors such as batch number, measurement variability, dropout events, and extreme measurements. PREFFECT offers several generative models (Table 1), each accommodating different types of data during the fitting procedure. All models require an unadjusted count matrix 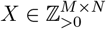 for the target tissue, where *M* is the number of samples and *N* is the number of transcripts, and can accommodate metadata (e.g., batch number, DV200, percent duplicates), and patient clinicopathological variables (e.g., grade, stage of a tumor). PREFFECT uses these variables to adjust the count data and for conditional inference [57] to improve the quality and interpretability of a fRNA-seq dataset.

**Table 1:** Types of PREFFECT models.

| Name | Tissues | Adjacency networks | Description |
| --- | --- | --- | --- |
| simple | only one target | no | Generative model can use sample metadata. |
| single | only one target | yes | GAT layer exploits sample-sample correlations. |
| full | multiple | yes | Borrows information across tissues. |

The structure of the *simple* model is similar to those used in scRNA-seq contexts, e.g., Lopez and colleagues [45]. The encoder produces an estimation *q*_Φ_ of the true posterior *p*_Θ_, where Φ and Θ are the sets of all relevant underlying parameters for *q* (i.e., weights in the neural network) and *p* (i.e., parameters of the underlying true distribution). *q*_Φ_ consists of several conditional VAEs including an (optional) encoder 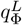, for the observed (log-)library sizes of the samples allowing the library size to vary during model fitting (Figure 3A, green).

**Figure 3.**
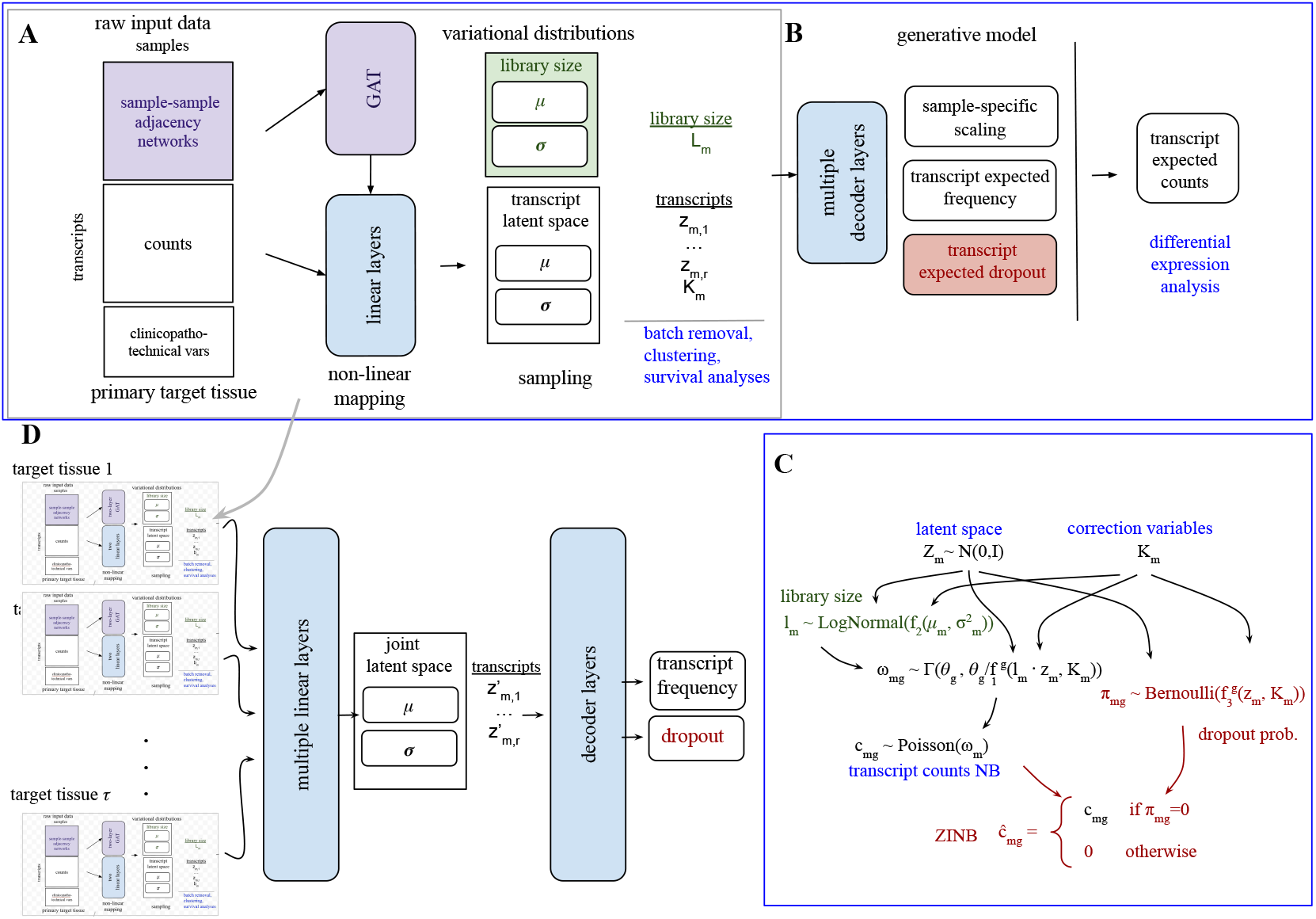
Overview of PREFFECT. **A**. The simple encoder consists of only the white boxes (count matrix and correction variables). The single encoder integrates the sample-sample adjacency graph with attention mechanisms (purple boxes). **B**. The single decoder extends the simple decoder with multiple decoder layers that allow integration of the adjacency information. If a ZINB is the desired distribution, the decoder also estimates a dropout rate *π* denoted in red. **C**. The relationships between all variables from the underlying statistical model. **D**. The full model combines one single encoder model for each available tissue.

**Figure 4.**
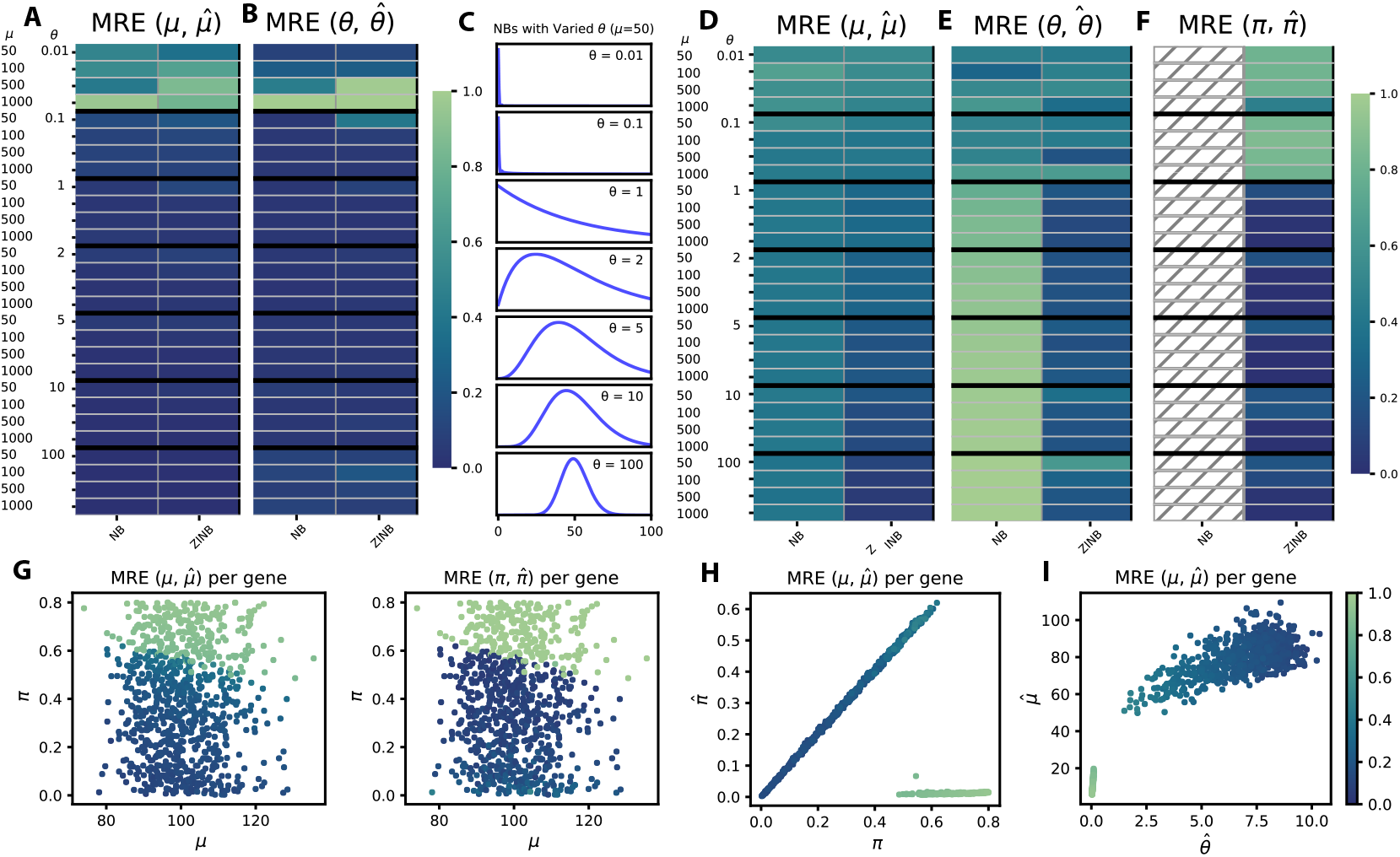
The ability of PREFFECT to recuperate generative parameters. NB counts were generated for *N* = 1, 000 transcripts across *M* = 1, 000 samples across a range of parameterizations for *µ* and *θ*. PREFFECT was then used to infer parameter levels under either an NB (left column) and ZINB (right column) model. Colors in the heatmap correspond to the mean relative error (MRE) between the generative parameters *µ* (**A**) and *θ* (**B**) versus their respective estimates 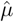 and 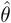. The MRE remains very low for all parameters except when *θ* is very small. **C**. Examples of the effect of *θ* on the NB distributions. When *θ* is very small at 0.01, many transcripts have a 0 count. **D, E, F**. The ability of PREFFECT to recuperate generative parameters when challenged with dropouts. The same synthetic generative methods from panels **A** and **B** are repeated but now each transcript was subjected to random dropout (from 0 to 80% of all samples are set to 0). PREFFECT was used to infer parameters. As expected, the ZINB model is near universally better than the NB model, where *µ* (**D**), *θ* (**E**) and *π* (**F**) have low MRE except for small dispersion *θ* levels. **G, H, I**. The performance of estimating ZINB parameters *µ* and *θ* with random amounts of dropout where *θ* = 10. **G**: Color is proportional to the MRE of the masked positions for each transcript (point) plotted according to the generative parameters *µ* and *π*. **H**: Color represents the MRE relative to *π* and 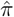 ; and **I**: MRE relative to the generative 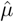 and 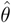.

The first decoder maps the latent spaces to fitted count matrices under an NB or ZINB distribution via three neural networks *f*_1_, *f*_2_, *f*_3_ (Figure 3C). Neural network *f*_1_ estimates the fraction of reads for each transcript in each sample *c*_*m,g*_ along with an inverse dispersion value *θ*. Neural network *f*_2_ estimates the library size *l*_*m*_ of each sample *m. θ* and *θ/l*_*m*_*· c*_*m,g*_ serve as the shape and rate parameter, respectively, to a Gamma distribution to form counts *ω*_*m,g*_ which serves as the rate parameter to a Poisson distribution and together define a NB distribution with mean *µ*_*m,g*_ and dispersion *θ* (see 4.1 for a more detailed explanation and also [45], Supplementary Note 3). If the user prefers to work with a zero-inflated model, a third neural network *f*_3_ estimates the logit of the dropout rate *π*_*m,g*_ for each transcript *g* in each sample *m. π*_*m,g*_ is used as the parameter of a Bernoulli distribution that models whether a transcript is observed or not. When combined with the NB distribution of *c*_*mg*_, this results in a ZINB distribution (Figure 3, red).

Loss is computed as a (possibly weighted) combination of the Kullback-Leibler (KL) divergence and reconstruction error (log-likelihood of the NB or ZINB as appropriate). A more detailed description of the neural network with details on dropout, choice of activation layers, injection of correction variables, and optimization is described in the Supplementary Methods 4.2.

### The simple model robustly estimates generative parameters when dropout rates are within observed ranges

We constructed synthetic count data that was generated in a manner that simulates the distributional properties of observed fRNA-seq datasets to explore the capacity of the simple model to recover generative parameters. Transcript count matrices were generated with a range values for the NB parameters *µ* and *θ* which captures the behavior of the vast majority (*>* 95%) of transcripts across the fRNA-seq compendium (Methods 2.3). Figure 4 shows that performance is near perfect for both *µ* (panel **A**) and *θ* (**B**) everywhere except when *θ* is very small (0.01). This corresponds to an NB distribution with almost all zeros (top of panel **C**), and is smaller than the value of *θ* for 99% of transcripts across the fRNA-seq compendium. Although the synthetic data was generated here using an NB distribution, the ZINB still accurately assessed the parameters *µ* and *θ* (second column of **A** and **B**).

The elevated number of zero counts for transcripts in fRNA-seq data motivated a study of how well PREFFECT can impute missing values. We used a simple self-learning approach to imputation where PREFFECT replaces masked values with the expected value from the estimated distribution. To explore this, we again used synthetic data generated with an NB distribution in the same parameter space described above. However, now each transcript is subjected to dropout with a randomly assigned rate *π ∈ U* (0 0.8), producing a ZINB distribution with known dropout locations.

Not surprisingly, the performance depicted in Figure 4D-E suggests the quality of the fits inferred by PREFFECT are overall poorer than simulations without dropout when the *NB* is used. However, when the ZINB is used, the performance remains high especially for larger *θ* values, and decreases only for low *θ*, likely because the presence of many endogenous zeros (zeros not caused by dropout) leads to an inflation in the estimate of 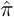.

To better investigate the role of the dropout rate *π* in parameter estimation, we constructed a new synthetic dataset where each transcript count is generated according to

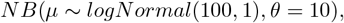

a choice of parameters that is unlikely to produce many endogenous zeros. If we mask values in this generated count matrix using a random dropout rate *π ∈ U* (0 .. 0.8) for each transcript, the estimate 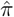 should be very close to *π*, since the vast majority of zeros are truly due to dropout. Figure 4G-I show that PREFFECT estimates *µ* and *π* well for *π <* 0.6 but degenerates for higher dropout rates. None of the datasets in our compendium had a dropout rate *≥* 0.55.

### Adjusting for batch effects

Especially in large-scale projects, technical limitations require that samples be prepared and profiled in batches. Batch-based profiling can introduce significant systematic effects that should be corrected for in order to ensure that downstream analyses are not unduly negatively affected. We explored the capacity of PREFFECT to identify and ablate batch effects.

In the first experiment, counts were generated for all samples using a family of NB distributions with location parameters determined by a single underlying transcript frequency vector *ω*. The samples were then randomly assigned to batch 0 or 1, but only counts for transcripts in batch 1 were systematically increased to simulate the batch effect (Methods 2.6). As expected, after training a simple PREFFECT model, samples clearly cluster by batch number when the frequency vectors are computed from the observed count matrix (UMAP, Figure 5A). However, by shifting samples in batch 1 towards batch 0 in the latent space, the resultant adjusted count matrices no longer cluster by their batch (Figure 5B). Figure 5C confirms that the frequency vectors computed from the simulation differ between batch 0 and 1, as expected. If PREFFECT successfully fits a good model, the difference in values between batch 0 in panel C and panel D will be marginal. The same statement holds for batch 1 between panels C and D. Lastly, by shifting batch 1 samples towards batch 0 in the latent space, batches 0 and 1 will have nearly identical frequency vectors, as observed in panel E, indicating a successful ablation of the batch effect.

**Figure 5.**
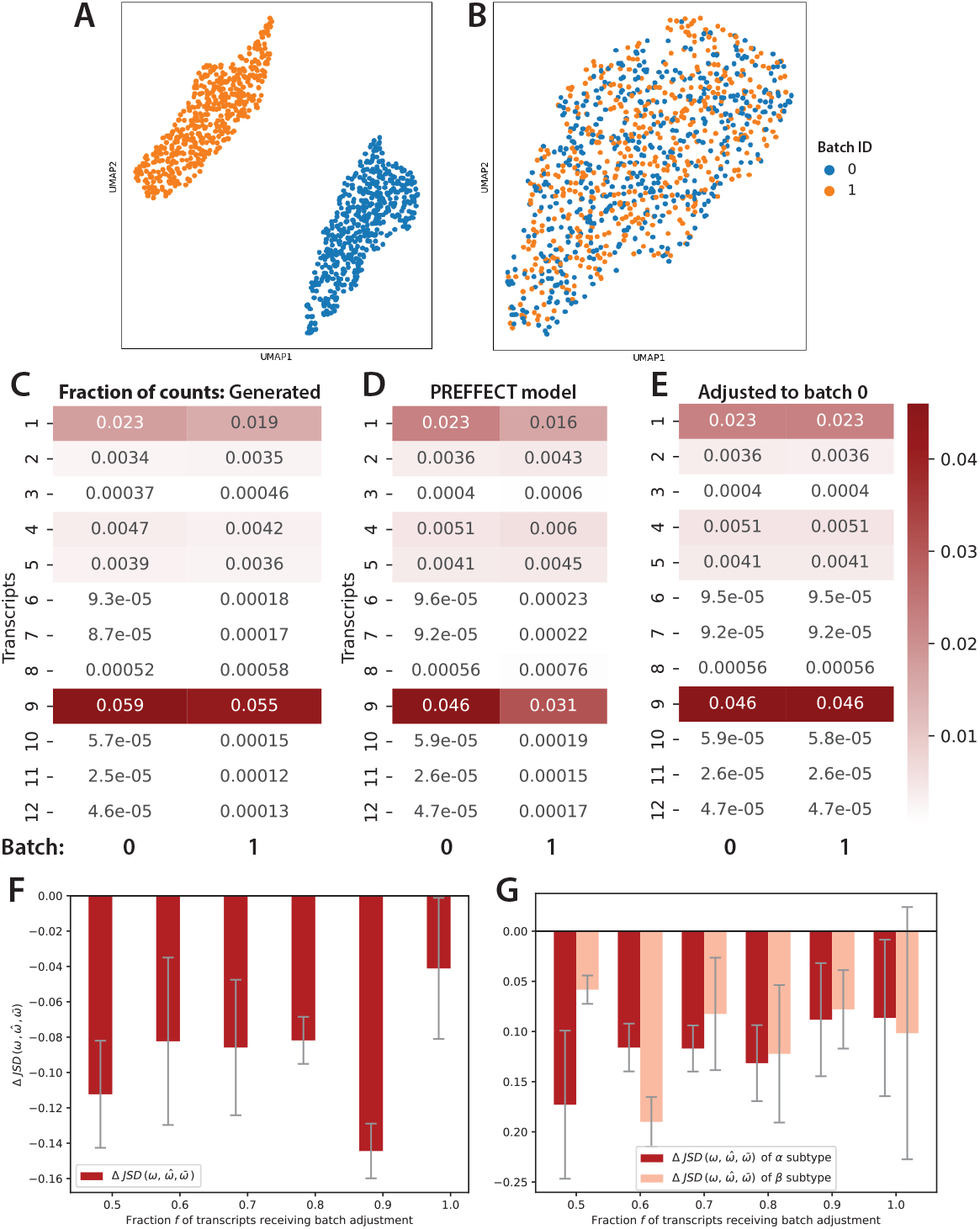
Adjusting for batch effects. A PREFFECT model was derived using synthetic data with a simulated batch effect on all transcripts with a randomly chosen subset of samples. **A**. UMAP embedding using the frequency vector *ω*_*s*_ for each sample *s* obtained from the raw count data. **B**. UMAP embedding using the estimated frequency vectors 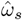 after adjusting the latent space of batch 1 to batch 0. **C**. The average fraction of all counts for each transcript between batches in the raw synthetic data. **D**. After training, these same fractions are retained when no adjustment is carried out. **E**. During inference, all transcripts were adjusted to batch 0. **F, G**. Correction with respect to fractional batch effect. Synthetic datasets were generated where counts for a fraction *f* of the transcripts (*f ∈ {* 0.5, 0.6, …, 1 *}*) in batch 1 were subjected to the effect. **F**. The difference in the Jensen-Shannon divergence (JSD) was computed between the true generative transcript frequencies *ω* and 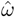, and between *ω* and the estimated frequencies after adjusting for the batch effect 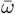. For all fractions *f*, this difference is negative implying that the adjustment has shifted the estimated frequencies closer to the true generative distributions. **G**. Similar to F but here samples were randomly assigned to either the *α* or *β* subtype with distinct transcript frequency vectors *ω*_*α*_ and *ω*_*β*_ respectively.

The second experiment tests the ability of PREFFECT to identify and adjust for a batch effect at different levels of pervasiveness. Here, a series of PREFFECT models were fit to a count matrix similar to the first experiment, but only a fraction *f* of the transcripts in batch 1 received the batch adjustment. We computed the difference between two similarities: (ii) the similarity between the generative transcript frequencies *ω* and the estimated frequencies 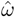, and (i) the similarity between *ω* and 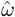 after adjusting for the batch effect with the latent space, denoted 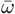. For all values of *f*, the batch adjustment 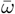 is more similar to the true generative *ω* (Figure 5F).

The third experiment depicted in Figure 5G extends the previous exploration to investigate cases where the samples may differ by both their batch and other effects, such as, for example, their subtype. To simulate this, samples were randomly assigned (with equal probability) one of two subtypes *α* and *β* each with a distinct frequency vector *ω*_*α*_, *ω*_*β*_ respectively. Again, regardless of the pervasiveness *f* of the batch effect, the adjustment reduces the degree of dissimilarity with the two true generative frequencies, although the improvements were somewhat reduced.

### Single tissue generative model: borrowing neighborhood information

Since fRNA-seq data has considerably high dropout rates and is prone to extreme measurements, we sought to incorporate additional information that could assist with the de-noising and imputation of count data. Toward this end, the so-called *single tissue* generative model incorporates a sample-sample correlation network represented as a *M × M* matrix *A*. Two samples are adjacency if and only if they are deemed sufficiently “similar”. Often similarity is defined by Pearson correlation distance or another metric applied to a subset of transcripts, but other techniques could be used either directly with the count matrix *X*, or some other independent datasets (not necessarily fRNA-seq), opening avenues for integrating multi-omics data, clinical information or other modalities.

In the single tissue generative model, the encoder processes the count matrix *X*, the adjacency matrix *A* alongside associated metadata *K* through a neural network to project the data into a latent space *Z*_*A*_. The top layer of this neural network uses a graph attention network (GAT) [47, 52], which takes as input transcript counts for each sample in addition to the sample-sample adjacency list (Figure 3A, purple). The attention mechanism helps the model focus on the most informative dependencies while minimizing the effect of others. GAT mechanisms also facilitate hierarchical feature learning by aggregating information over multiple layers and at different levels of granularity. As before, the (log) library sizes (*log L*) can be allowed to vary during training. The encoder produces an estimation *q*_Φ_(*Z*_*A*_, *Z*_*l*_ |*X, A, log L, K*) of the true posterior distribution *p*_Θ_, where Φ consists of all parameters of the neural network and Θ represents the true parameters underlying the data. Linear layers are applied after the GAT to form the final latent encoding *Z*_*A*_ of the graph and count matrix. The decoder reconstructs an estimate of the count matrix *X* and the adjacency matrix *A*.

### The single tissue generative model improves sample clustering

We explored how the inclusion of the adjacency matrix could improve performance of downstream tasks, specifically sample clustering. Since available fRNA-seq datasets are limited in size, we generated a pseudo-synthetic dataset consisting of 200 samples for each of the 5 breast cancer subtypes (Methods 2.4). Briefly, a count matrix was generated which contains the PAM50 transcripts estimated from breast cancer datasets in the compendium. The frequency vectors for the five subtypes depicted in Figure 6A are in broad concordance with the original PAM50 frequency vectors (Figure 2A). Edges were included in the adjacency matrix if and only two samples had the same subtype.

**Figure 6.**
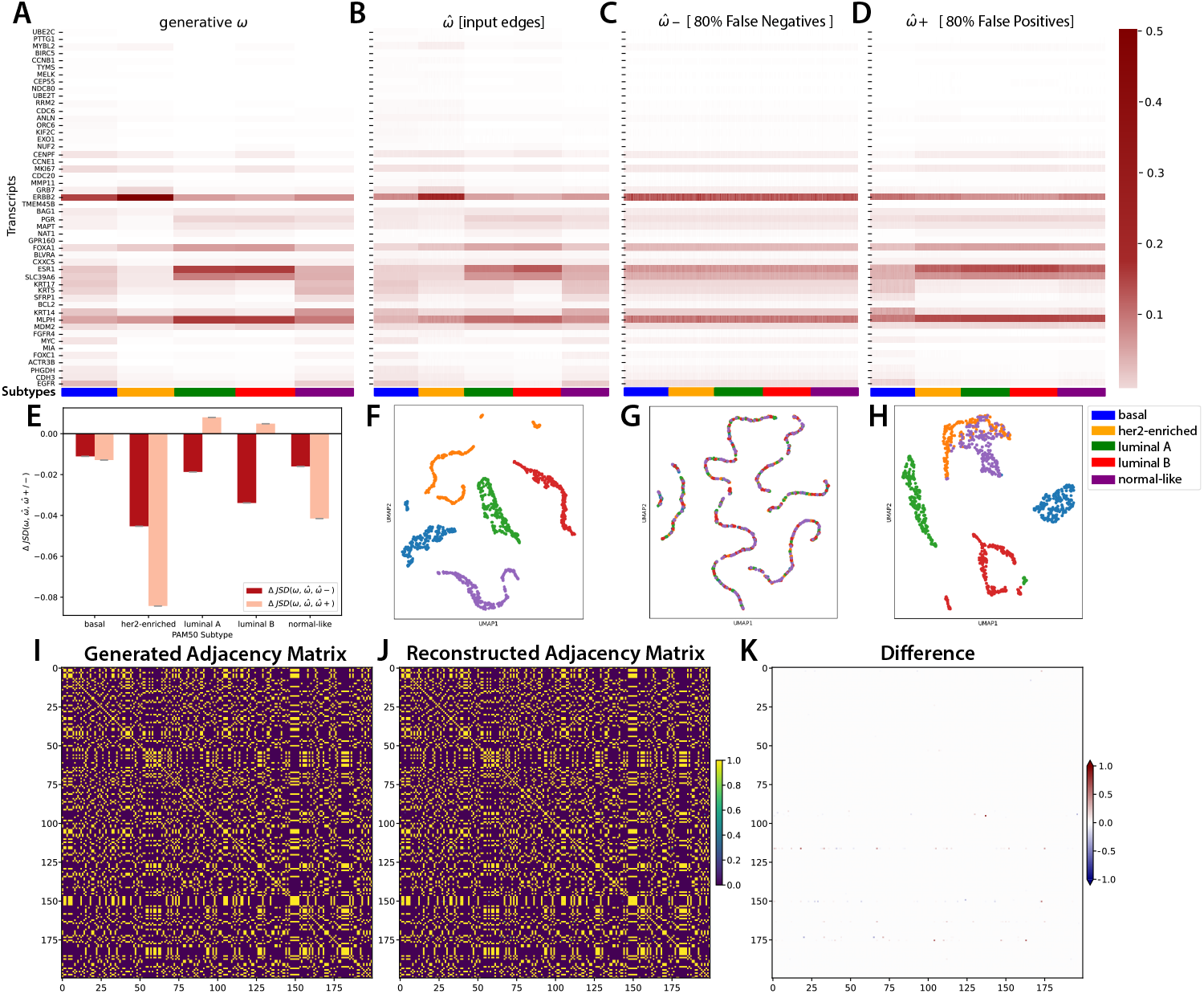
The adjacency information assists in down-stream applications such as sample clustering. A pseudo-synthetic count matrix was constructed using NB distributions for the PAM50 transcripts derived from breast cancer fRNA-seq datasets. **A**. A heatmap of frequency vectors *ω* for each of the 5 PAM50 breast cancer subtypes. **B**. A heatmap inferred 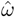 of a PREFFECT model built using an informative sample-sample edge matrix. The UMAP is also clustered from the 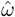. **C**. The 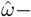 when 80% of edges of the sample-sample adjacency matrix were randomly deactivated. **D**. The 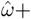 when 80% of edges of the sample-sample adjacency matrix were randomly activated. **E**. We compare the distributions of 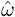 from **B-D** to the generative *ω* (**A**) of each subtype via JSD. We plot Δ *JSD* of 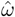 (**B**) to either 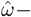 or 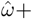 (**C-D**) across each PAM50 subtype. **F-H**. UMAP clustering of 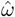 from **B-D**, respectively. **I**. The adjacency matrix for a random subset (*M* = 200) of the samples. Here yellow corresponds to the existence of an edge indicating both samples have the same subtype (probability of 1) and black corresponds to no edge (probability of 0). **J**. The reconstructed adjacency matrix. **K**. Differences between the generated and reconstructed adjacency matrices are negligible.

Not surprisingly, the single-tissue model trained with both the count data and the adjacency matrix was able to recuperate the subtype-specific transcript frequency vectors *ω* (average JSD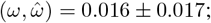 Figure 6B). Moreover, the UMAP produces five distinct clusters that nearly perfectly separate samples by subtype (panel F). To test the contribution of the adjacency network, we repeated the training process but this time removed a fraction of all edges. Figure 6C depicts the estimated frequency vectors when 80% of all edges were removed, leaving only 20% of the edges between samples of the same subtype. There is a noticeable decrease in the model’s capacity to recapitulate the generative transcript frequencies *ω* (average 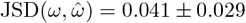). This is reflected in the associated UMAP where each “snake-like” cluster contains samples with different subtypes (panel G). The last experiment instead introduces false positives into the adjacency matrix prior to training. Again, the capacity of the model to recapitulate the *ω* vectors is reduced (average 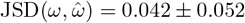; Figure 6D) and the resolution of the UMAP has decreased with some her2-enriched samples clustering with normal-like samples, and some confusion between luminal A and B (panel H). Although reconstruction of the adjacency matrix required unequal weighting of the reconstruction and KL-associated loss (Methods 2.7), the reconstructed graphs were nearly indistinguishable from the original input graphs (Figure 6I-K).

### Full model: Incorporating surrogate tissues

In some cases, surrogate fRNA-seq data is available for two or more related tissues or conditions. For example, in disease studies, often both the affected and matched healthy/normal tissue from a patient is profiled. In cancer, the profiles of an index lesion can be complemented by profiles of their match normal tissue, the tumor microenvironment, and metastatic sites. The full model in PREFFECT generalizes the single model by accepting a set of tissue-specific count matrices 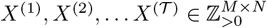 along with corresponding sample-sample adjacency networks *A*^(1)^, *A*^(2)^, … *A*^(^ *𝒯* ^)^ for the *𝒯* matched tissues profiled over a common set of *N* transcripts (Figure 3D). The learning procedure fits the NB or ZINB models to each tissue via the GAT layers and adjacency matrices as in the single layer model, but additional layers in the network combine the *𝒯* latent spaces into a single joint latent space. In this way, the related tissue types influence the fit (Figure 7).

**Figure 7.**
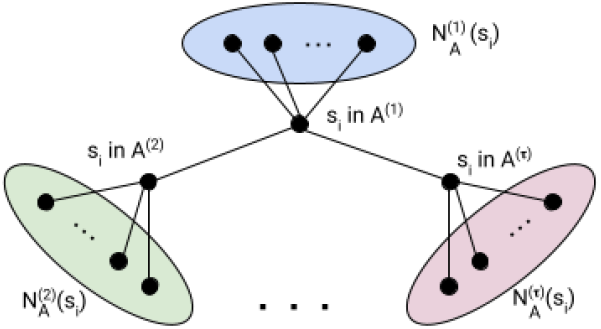
Neighborhood information is encoded in the adjacency networks. Blue ellipse indicates the neighborhood of a specific sample *s*_*i*_ in the target tissue, while the green and pink ellipses depict the neighborhoods of sample *s*_*i*_ in the first and second surrogate tissues. Edges are included between nodes corresponding to *s*_*i*_ between the three tissues.

### Profiles of matched tissues can improve sample clustering

We again use breast cancer as an example, since we have access to a dataset with matched primary tumor and stroma profiles. It is well-established that breast cancer samples have both a primary tumor subtype [55, 58] and a tumor stroma subtype (e.g., [59]). The tumor and stroma subtyping schemes are distinct and the relationship between them appears to be complex and is still not fully understood. The full PREFFECT model provides a means to generate clusterings informed by the transcriptional profiles of both tissues simultaneously. To explore this, we generated pseudo-synthetic count matrices for both tissues as follows (Methods 2.9). Starting with the primary tumor matrix, 1000 samples were randomly assigned to four breast cancer PAM50 subtypes; each sample had counts for 100 transcripts corresponding to the PAM50 transcripts and 50 stroma-specific transcripts. For the tumor count matrix, the PAM50 transcripts were generated randomly according to an NB distribution with parameters estimated from breast cancer fRNA-seq datasets in the compendium. However, the 50 stroma-specific transcripts were assigned random counts according to an NB distribution ignoring all subtype information (that is, these transcripts are uninformative in the tumor profiles). For the stroma count matrix, each samples was also assigned a stromal subtype: basal samples were randomly assigned either stromal subtype *α* or *β*, her2-enriched samples were assigned stromal subtype *γ*, and luminal A and B samples were assigned stromal subtype *δ*. A distinct frequency vector was generated for each stromal subtype and used to generate counts for the 50 stromal genes. Random values were assigned to the PAM50 transcripts in the secondary tissue dataset (that is, the PAM50 transcripts are not informative in the stromal samples). In this two-tissue scenario, we would expect that samples will cluster according to the five tumor subtypes when the tumor count matrix is used, but *α* and *β* samples would not separate. Conversely, we expect that the samples will cluster by the five stromal subtypes when the stromal count matrix is used, but the luminal A and B samples would not separate.

Figure 8A confirms this hypothesis regarding separation of the luminal A and B subtypes when using the tumor counts. Interestingly, in the stroma-related Figure 8B, we also observe the luminal A and B samples separated. We hypothesize that this unexpected result is due to backpropagation during model training, which transfers information from the model of tumor counts to the model of stromal counts. Regardless, when the combined latent space is used to cluster the samples, both the tumor luminal A and B subtypes and the stromal *α* and *β* subtypes are separated (Figure 8C).

**Figure 8.**
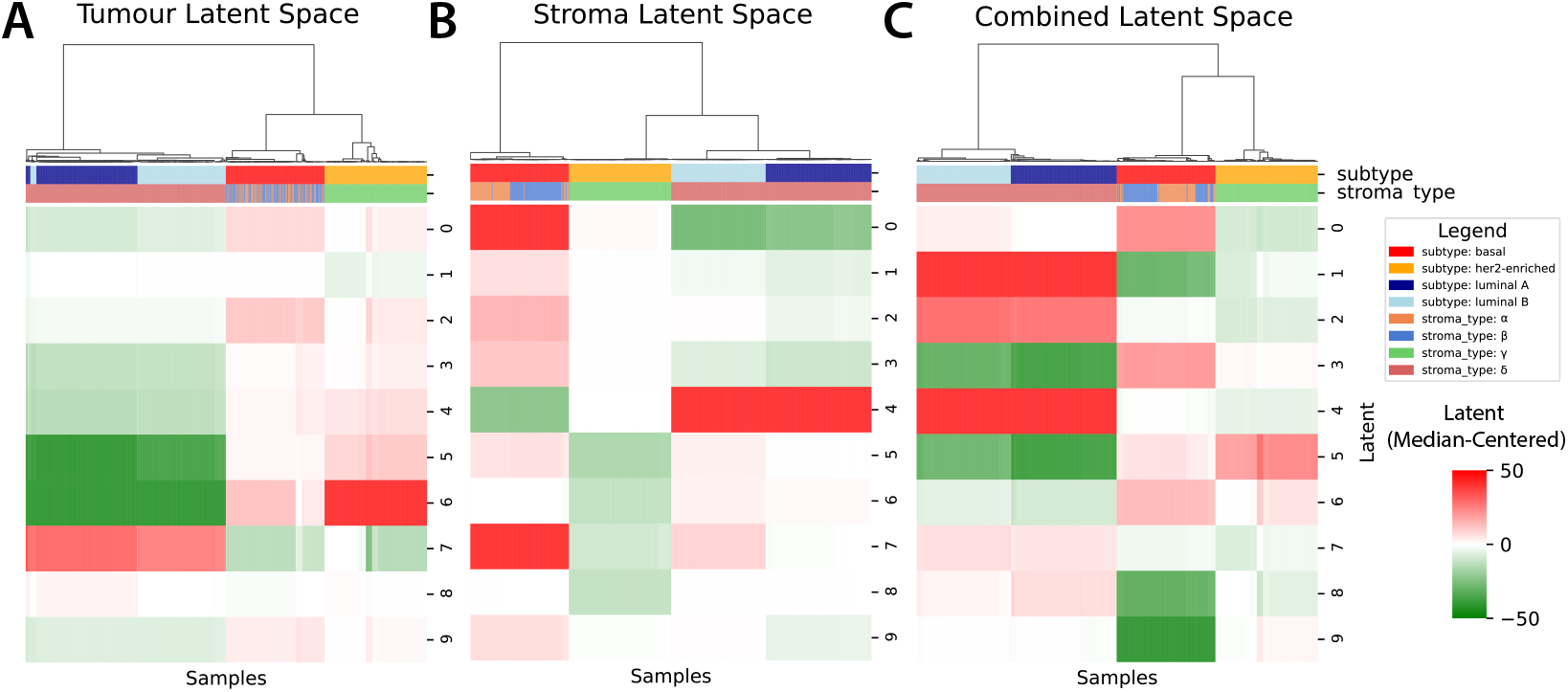
Multi-tissue PREFFECT models can be developed using information from both primary and secondary matched tissues. To test the full model, we developed both a pseudo-synthetic breast cancer tumor count matrix (where PAM50 transcript counts follow subtype-specific distributions) and a secondary synthetic stromal count matrix (where a second set of 50 transcripts had counts separating them into four stromal subtypes *α, β, γ, δ*). Hierarchical clustering was applied to the resultant latent spaces **A** The latent space of the primary tumor tissue, which clusters according to PAM50 subtype but not stromal subtypes. **B** The latent space of the secondary stromal tissue, which clusters according to stromal subtype but also surprisingly separates luminal A from B in the tumor, likely due to information transfer during the learning procedure. **C** The combined latent space, which clusters the samples by the cross-product of the two subtyping schemes.

### PREFFECT models for contemporary fRNA-seq datasets

We examined the capacity of PREFFECT to fit good models with available fRNA-seq datasets, and tested whether the resultant models aided in downstream analysis, specifically sample clustering. Simple models were built for each dataset in the compendium, but our analysis below focuses once again on the six breast cancer-related datasets to investigate performance. Here, models were restricted to *N* = 776 genes from the well-studied pan-breast cancer BC360 panel (NanoString Inc.), since we can expect that their transcript values will vary significantly across the datasets. Initially, hierarchical clustering was applied to the data before applying PREFFECT (log-transform mean trimming 1% with variance stabilizing transformation). We observe a broad range of count values with many zero counts (represented by white). Although clusters are enriched for same-subtype samples (especially basal), many subtypes (especially luminal B) are diffuse across the clustering (Figure 9A for dataset *GSE167977*). When the inferred transcript frequencies 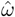 are used instead, the expression becomes more polarized away from 0, presumably due to imputation of missing values (panel B). When the samples and transcripts are reclustered using the inferred transcript frequencies, we see much more homogeneous clusters for every subtype with the exception of the basal subtype which was already homogeneous.

**Figure 9.**
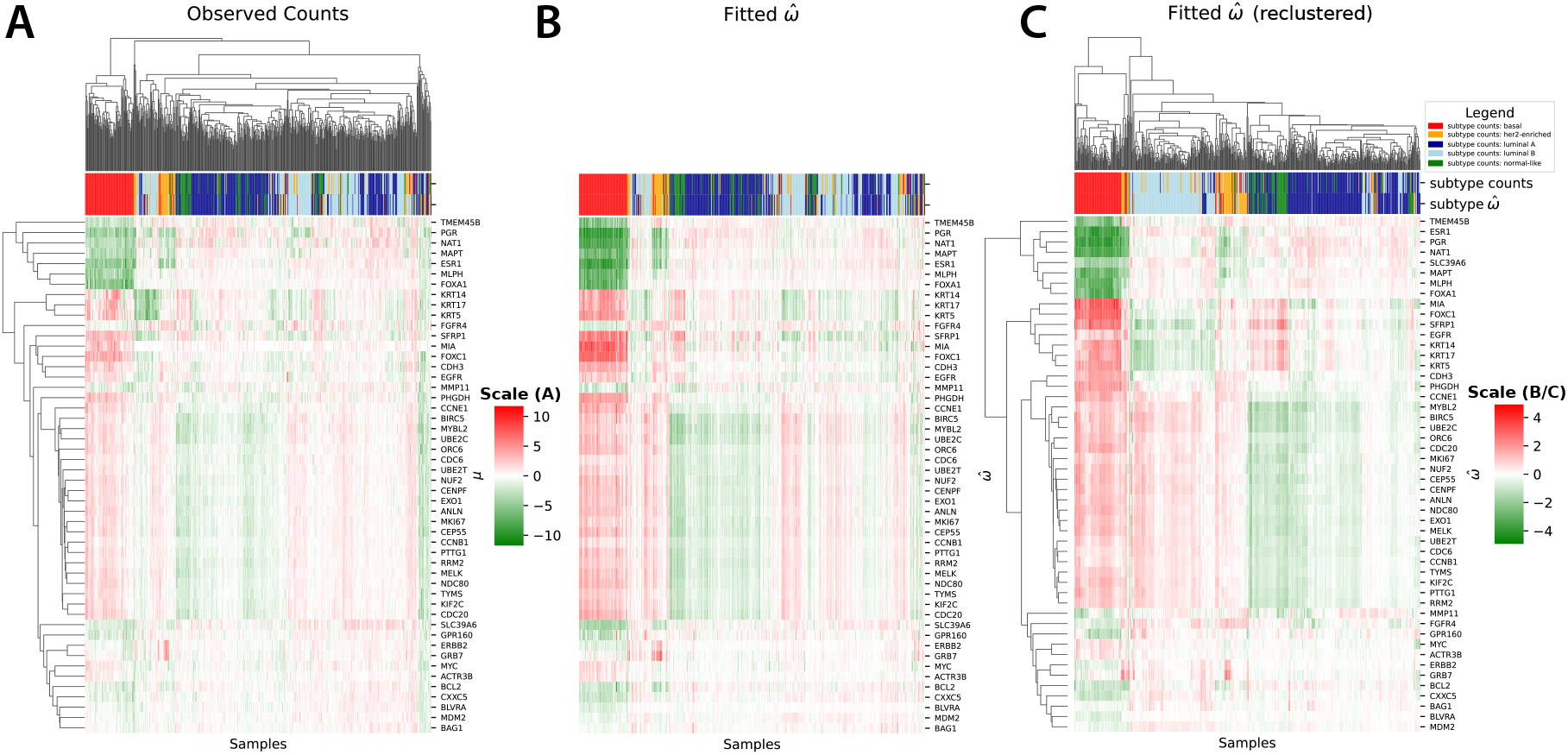
A simple PREFFECT models was fit using the BC360 (NanoString Inc.) panel from dataset *GSE167977*. **A**. Hierarchical clustering was performed using the PAM50 transcripts contained in the BC360 panel. **B**. The same sample and transcript clustering is re-drawn from panel *A* but instead here color corresponds to the estimated transcript frequencies 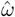. **C**. Using the estimated frequencies 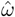, the samples and transcripts are re-clustered, resulting in sample clusters which are more homogeneous, consisting of a single subtype. In all panels, PAM50 subtypes were inferred from observed counts (top) and also from the inferred frequencies 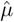 (bottom).

## Discussion

Analysis of the fRNA-seq compendium suggests that the vast majority of transcript counts are well-modeled by an NB distribution, although both the mean *µ* and dispersion *θ* parameters vary considerably compared to other types of RNA-seq data including bulk and single-cell profiling. Many transcripts have a considerable number of zero counts; in fact, the median zero count rate in all datasets was almost half (0.46). Nevertheless, the NB is still able to model transcript counts well with small *µ* and small *θ <* 1 indicative of an “exponential-like” shape. Few transcripts were better modeled using the ZINB with its additional dropout parameter *π*, an observation that holds across all datasets. Although the distribution of counts for many transcripts is suggestive of a mixture of a zero-inflated Poisson and truncated normal distribution [32, 33], a simple NB distribution was still better able to fit the data, most likely because their variances are far greater than their means.

We focused here on breast cancer datasets and often specifically on subtype in our analyses. This restricted focus allows us to comment on the capacity of fRNA-seq profiles to reflect the well-established properties of these samples. For example, our investigation of the PAM50 transcripts highlighted large differences in the shape of the NB distribution between the subtypes. Her2-enriched samples, which we expect to highly express the *ERBB2/HER2* gene, do in fact have counts that fit well to a NB with a large *µ >* 6, variance and dispersion. In contrast, non-her2-enriched samples have a significantly lower *µ* (decreased 6 to 12-fold) and a greatly reduced variance (up to a 506-fold differential), with a distribution shape similar to an exponential. A similar statement can be made for the majority of the PAM50 transcripts. Not surprisingly, the count matrices from essentially all datasets in the compendium were affected by batch effects and other technical variables, including DV200, which measures the average length of RNA fragments.

In response, we created PREFFECT, a series of generative models based on conditional VAEs to impute and factorize observed transcript count data to de-noise and adjust for both technical and biological variation. PREFFECT offers a number of alternatives to adjust for batch effect, which is a difficult problem likely without a single unique solution. We continue to experiment with alternative approaches based on disentanglement [60] and by incorporating adversarial learning to “cleanse” the latent space of batch information [61, 62]. Such approaches may better remove the effects of batch or other user-specified adjustment variables.

Using so-called pseudo-synthetic datasets of sufficient size, we show that PREFFECT can accurately infer generative parameters and accurately impute missing values for the range of parameter values observed in the real data. The high frequency of zero counts in the fRNA-seq data motivated extensions of the simple model that integrates neighborhood information via a sample-sample adjacency graph. This single tissue model uses graph attention networks (GATs) to assist the learner to attend to the most influential neighbors from which to infer missing values. We showed how such imputation can lead to better patient subtyping with breast cancer datasets. The full model allows for multiple matched tissues from the same patient sample to be integrated. To the best of our knowledge, this is the first generative tool to incorporate multiple patient-matched tissues. We show that the joint latent space that PREFFECT forms can lead to more insightful patient clustering, again in the context of breast cancer subtype.

The vast majority of available fRNA-seq datasets currently are of moderate size. For example, the datasets in the compendium used here have a median of 93 samples, a value that is four orders of magnitude smaller than some datasets available for scRNA-seq. Generative approaches such as PREFFECT, which provide a means to ablate nuisance technical parameters and better capture true biological signal, would certainly benefit from larger fRNA-seq datasets, given the degree of variability and extreme measurements, especially in contexts such as cancer where we know that samples are affected by strong transcriptional programs (e.g., estrogen receptor status in breast cancer and other subtype-related programs). The undersized nature of current fRNA-seq datasets may partially explain why FFPE-based studies remain very challenging. Nevertheless, PREFFECT is able to fit models to existing datasets and we are able to show that the models may assist in improving downstream analyses. However, for some datasets, this required multiple training runs with different parameter settings that appear dataset specific. Model fitting to larger fRNA-seq datasets would likely be more straightforward.

PREFFECT would benefit from various extensions including support for differential expression with frameworks such as Boyeau et al. [63] and more support for visualization of the various latent spaces and fitted data. The full model with multiple tissues is complex and its success still requires rather extensive experimentation at times with multiple parameters to find good weightings for components of the loss function and other parameters involved in training. We are continuing to simplify and develop best-practice recommendations for integrating multiple tissues.

## Methods

`PREFFECT` is implemented in `Python` version 3.9 [64] with PyTorch version 1.12, the `geometric` package version 2.3 [65] and the `CUDA` toolkit version 10.2.89. Statistical analyses were performed using `Python` version 3.9 and `R` version 4.4 [66]. Hierarchical clustering was performed using the `seaborn clustermap` function with average distance linkage and Euclidean distance. UMAP was performed using the `umap` *−*`learn` [67] package in SCANPY [68] (`min_dist = 0.3`).

### 2.1 The fRNA-seq compendium

fRNA-seq datasets were downloaded and processed, only when raw count information was available. All datasets with GSE identifiers were obtained from the Gene Expression Omnibus https://www.ncbi.nlm.nih.gov/geo/. The TMBC dataset was obtained from The Metastatic Breast Cancer Project [54] available through cBioPortal [69]. The Sunnybrook cohorts are not currently publicly available but are pending publication. We collected clinicopathological and technical variables (such as batch identification number) whenever possible. The datasets vary across tissue, cell type, age of the cohort and profiling techniques (Supplementary Table S1). Transcripts with zero counts across all samples were removed. Since the PAM50 subtype for samples was not provided in the breast cancer FFPE datasets, we estimated it using the PAM50 classifier [55]. In cases where a PREFFECT model was not used, the data was first subjected to a variance stability transformation, log-transformed and gene-centered. The resultant counts were to centroids for each of the PAM50 subtypes (obtained from `genefu` version 2.22.1) using Pearson correlation. The molecular subtype with the highest concordance was selected for each sample.

### 2.2 Fitting statistical models to fRNA-seq datasets

The following methods apply only to our initial descriptive statistics investigating the fit of different distributions to fRNA-seq data. To mitigate differences in the number of reads per sample, each input count matrix 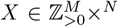, where *M* and *N* are the number of samples and transcripts respectively, was transformed to matrix *X* where

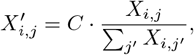

for a large constant *C* (e.g., *C* = 10^6^). Where noted, we used a trimmed mean approach to mitigate the influence of outliers (set to 1%), although this was not used when comparing the NB versus ZINB distributions (Figure 1B), as increasing the trimmed mean percentage increases the number of zeros that are removed and therefore could result in bias against zero-inflated distributions such as the ZINB.

Expression data was fit to six distributions using two different methods. The AIC measures the goodness of fit as 2*k −* 2*ln*(*L*) where *k* is the number of parameters of the distribution and *L* is the likelihood of the model given the observed data. All non-zero inflated distributions were fitted using the `StatsModels` package [70]. For zero-inflated models, the AIC and distributional parameters were estimated using the `minimize` function from scipy [71] to optimize the negative log-likelihood; this was repeated over a range of initial *π* values to minimize the negative log-likelihood. The *D* statistic of the Kolmogorov–Smirnov (KS) test was computed between the empirical cumulative distribution function for each transcript in every dataset and the cumulative distribution function of each reference distribution. The KS test was used to ensure that differences in the number of parameters between the six distributions did not unduly affect the results. In this analysis, transcripts were removed if they did not have non-zero counts in at least 20% of the samples. A 1% trimmed means was applied to eliminate outlier measurements.

### 2.3 Synthetic datasets

Synthetic datasets were generated primarily as a convenient means to investigate technical correctness, the limits of the PREFFECT generative models and some parameter optimization. For the analysis related to Figure 4**A-F**, count matrices were generated across a range of location and scale values for the NB distribution. For each pair (*µ, θ*) *∈ {* 50, 100, 500, 1000*}× {*0.01, 0.1, 1, 2, 5, 10, 100 *}*, a count matrix was formed across the *N* = 1000 transcripts and *M* = 1000 samples using variates from *NB*(*µ, θ*). For the analysis related to Figure 4**G-I**, the count *R*_*g*_ for each transcript *g* in each sample is formed as follows:

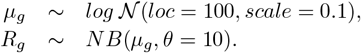

In several places, we require count data generated by a ZINB process. Toward this end, each transcript in the dataset was assigned a dropout value *π ∼ U* (0.0.. 0.8). Then *π* samples across the transcript were randomly selected and these values were set to 0. Batch effects were simulated as follows: (1) samples were included in batch 0 with probability *α* (sampled from *U* (0, 1)), otherwise they were assigned to batch 1. (2) Two location parameters *µ*_0_ and*µ*_1_ are chosen for the batch-specific NB distributions. (3) For each transcript *i* in sample *j* belong to batch *k, µ*_*i,j*_ *∼ logNormal*(*µ*_*k*_, 1) and *R*_*i,j*_ *∼ NB*(*µ*_*i,j*_, *θ*). Different values for *µ*_0_, *µ*_1_ and *θ* were explored.

### 2.4 Pseudo-synthetic datasets

Since the fRNA-seq datasets are small (*<* 200 samples), we used a pseudo-synthetic approach in our studies of the single tissue generative model and sample clustering. Here we fit an NB distribution to the counts for each PAM50 transcript, obtained from real datasets and stratified by patient subtype after 1% mean trimming. Patient subtype was estimated using the PAM50 classifier [55] after applying a variance stabilizing transformation to the counts and log-transforming the data. The patient subtype was estimated again after fitting the PREFFECT model without this transformation. Additional transcripts were added to the matrix but here counts were simply random variates following the methods described in Methods 2.3. When required, a sample-sample adjacency matrix was constructed by placing an edge between two samples of the same subtype. To evaluate the contribution of the adjacency matrix to imputation, we constructed a null adjacency matrix by randomly permuting the edges in the matrix. We ensured as best possible that the samples had similar library sizes regardless of their subtype.

### 2.5 Imputation

Several experiments were carried out to test the capacity of PREFFECT to self-learn or *impute* missing values using synthetic or pseudo-synthetic datasets. The collection *D* of positions (*i, j*) in the count matrix that were replaced by a zero to simulate dropout was recorded. The median relative error was computed between the true (hidden) generated and the imputed value across all such locations (*i, j*): For *X ∈ {µ, θ, π}*,

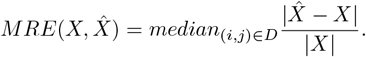

### 2.6 Batch corrections

Several experiments were carried out to investigate the capacity of PREFFECT to adjust for batch effects. In the first experiment, we created a synthetic dataset with *M* = 1000 samples and *N* = 900 transcripts and simulated a simple batch effect as follows. First, a sample *s* was assigned to batch 0 with probability *b*. Otherwise, it was assigned to batch 1. Let *batch*(*s*) denote its batch. Second, a frequency vector *ω* for the transcripts was generated using the stick-breaking algorithm [72]; *ω* was used to generate variates for all samples, regardless of batch and a single suitably large library size *L* was chosen for all samples. Transcript counts *C*_*s,g*_ for sample *s* and transcript *g* were generated according to a hierarchical model as follows:

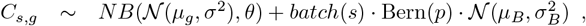

where *µ*_*g*_ = *L · ω*_*g*_, *σ* = 1, *p* corresponds to the probability a transcript is subject to the batch effect, *µ*_*B*_ and 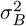 are the parameters for a normal distribution describing the batch shift. Note that the final library sizes (from summing over all transcripts per sample) will tend to be larger for batch 1 samples.

Three distinct experiments were conducted as follows:

*Experiment* 1. Library size *L* = 10^5^; *B* = 100; 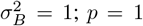 implying all transcripts in batch 1 received the adjustment. Samples were assigned to the two batches with equal probability.

*Experiment* 2. As for Experiment 1 but the probability *f* that an individual transcript in batch 1 would be subjected to the batch effect varied from 0.5 to 1.

*Experiment* 3. Similar to Experiment 2 but two distinct frequency vectors *ω*_*α*_ and *ω*_*β*_ were randomly generated via the stick-breaking algorithm representing two subtypes *α* and *β* respectively. A sample was randomly assigned to either subtype *α* or *β* with equal probability. Therefore, a transcript *g* in sample *s, µ*_*g*_ is equal to either *L · ω*_*α*_ or *L · ω*_*β*_ depending on whether *s* was assigned to subtype *α* or *β*.

The *Jensen–Shannon divergence*, a symmetrized and smoothed version of the KL divergence, is used to measure the distance between frequency distributions the generative transcript frequencies *ω* and the estimated transcript frequencies 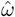. The JSD is a symmetrized and smoothed version of the KL divergence:

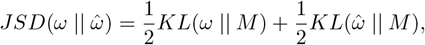

where 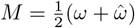) is the average, mixture distribution and KL(*·, ·*) is the KL divergence. In Figure 5F and G, we are interested in measuring the change in JSD after adjusting for the batch. In particular, we compute

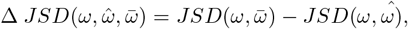

where 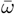 is the adjustment (via the latent representation) of 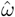.

Using the estimated transcript frequency vectors *ω*, the samples were mapped to two dimensions for visualization using UMAP.

### 2.7 Construction of adjacency matrices

The single and full models accept as input a sample-sample adjacency matrix *A*^(*τ*)^ for each tissue *τ*. If the adjacency matrix is formed from count data, which is not necessarily the same as the input matrix *X*^(*τ*)^, it is recommended that it be first transformed (e.g. square root and mean trimming) and normalized. In the experiments related to clustering with the single and full models, edges were included in the graph if and only if the two samples were of the same subtype.

### 2.8 Investigating the contribution of the adjacency matrices

Our explorations of the contribution of the sample-sample adjacency network to imputation, clustering and network reconstruction, used the breast cancer datasets from the compendium. In particular, we needed to generate a (large) set of pseudo-synthetic samples using a transcript frequency vector *ω*_*s*_ specific to each subtype *s*. We began by preparing each dataset as per Methods 2.2 before PAM50 subtype was estimated for each sample. Next, we transformed the raw input count matrix *X* to a matrix *X*″ where

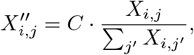

but this time we used *C* = 1 so that the resulting rows (samples) of *X*″ correspond to the frequency vector for all transcripts. Now for each subtype *s*, we compute a vector *ω*_*s*_ which is the average frequency of each transcript in all samples of the subtype *s. ω*_*s*_ is then used to generate sample-specific variations across each of the genes. For each subtype *s*, a chosen library size *L*, and gene specific dispersion *θ*, we create *N*_*s*_ pseudo-synthetic samples where each such sample has a transcript count vector formed as follows:

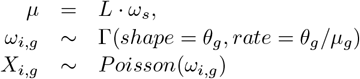

As such, each transcript should be distributed according to a *NB* distribution with parameters mean *µ* and dispersion *θ* as per the generative model underlying PREFFECT. An adjacency matrix was formed as per Methods 2.7 with edges placed between two samples if and only if they are of the same subtype.

Successful training of the model required a specific weighting scheme to balance reconstruction versus KL-associated loss. After a search across a broad range of possible values, we found that edge reconstruction was most accurate when the components of the loss function were waited as follows: KL weight adjusted by a factor 0.1, expression reconstruction adjusted by a factor of 100, and edge reconstruction weight by 1000.

### 2.9 Investigating the full model

This experiment requires two matched count matrices; the first matrix representing the primary tumor transcript counts and the second matrix representing the stromal counts. In both cases, there are 100 transcripts and 1000 samples where sample *i* in the primary tumor matrix is matched with sample *i* in the stromal matrix. For the primary tumor count matrix, assigned 250 (of 1000) samples to each of four breast cancer subtypes (here we did not use a normal-like subtype for simplicity). The first 50 transcripts correspond to PAM50 genes. NB counts were generated from parameters estimated for each subtype from breast cancer datasets in the fRNA-seq compendium. The second 50 transcripts are non-informative; the counts are simply variates from NB(100, 1) and so should have no significant impact on downstream sample clusterings. This gives us a set of *M* = 1000 samples each assigned a tumor subtype across the 50 PAM50 transcripts and 50 non-informative transcripts.

To simulate stromal subtyping schemes, we partitioned the 1000 samples into 4 stromal subtypes (labelled *α − δ*) as follows:

- if the sample was assigned the basal subtype in the tumor samples, it randomly receives either stromal subtype *α* or *β* with equal probability;
- tumor her2-enriched samples were assigned stromal subtype *γ*; and
- tumor luminal A and B samples were both assigned stromal subtypes *δ*.

In this (artificial) manner used for exemplary purposes, the stromal subtypes provide greater refinement of the tumor subtypes in one case (basal subtype), and the tumor subtypes provide greater refinement of the stromal subtype in one case (the *δ* subtype is fractured into luminal A and B subtypes).

A distinct transcript frequency vector was generated via a stick-breaking algorithm for each of the 4 stromal subtypes (as per Methods 2.6). The stromal count matrix consists of the 50 PAM50 transcripts and 50 stroma-specific transcripts. For the stroma, the PAM50 transcript counts are just random variates from NB(100, 1). The 50 stroma-specific transcript counts are generated using the corresponding frequency vectors *ω*_*α*_, *ω*_*β*_, *ω*_*γ*_, *ω*_*δ*_.

### 2.10 Optimization

Supplementary Information 3 provides a detailed description of the model and loss functions. The Adam optimizer was used for all hyper-parameter searches that included various learning rates, weight decay, number of epochs, mini-batch size, size of the intermediate *r*′, size of latent space *r* (*r* also controls the number of heads in the GAT layer), and the total number of samples *M*. The *α* parameter for leaky ReLU activation functions was set to 0.2 and dropout was set to throughout all analyses.

## CODE AND DATA AVAILABILITY

PREFFECT is available at https://github.com/hallettmiket/preffect. The code used for analyses in this paper is available at https://github.com/hallettmiket/preffect-paper.

## 3 Supplementary Information

**Supplementary Table 1:**
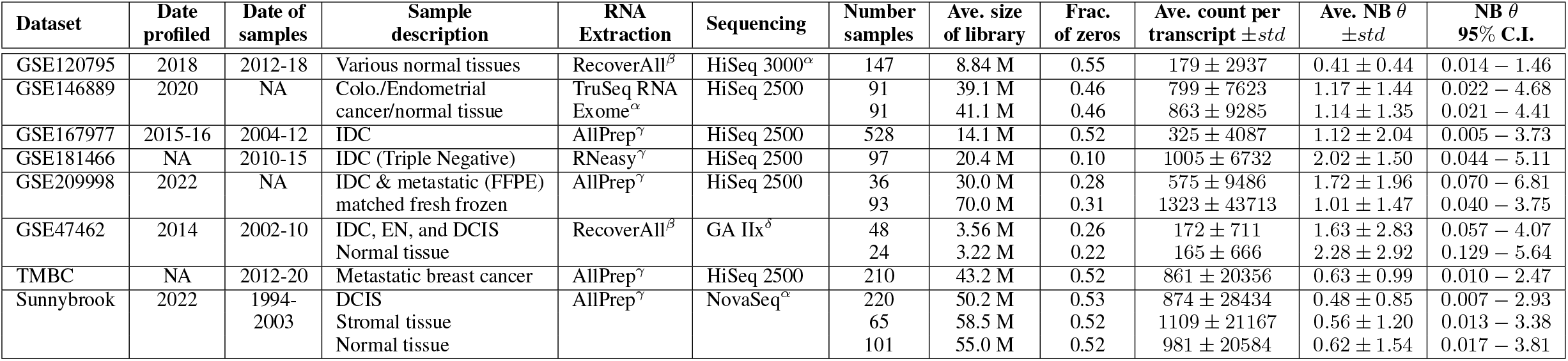
The fRNA-seq compendium. IDC: Invasive Ductal Carcinoma of the breast; EN: Early Neoplasia of the breast; DCIS: Ductal Carcinoma *in situ*; NA: Not Available; *α*: Illumina Inc. *β*: Ambion/Life Technologies Inc. *γ*: Qiagen Inc. *δ*: GA IIx: Genome Analyzer IIx, Illumina Inc.

### AIC versus Kolomogorov-Smirnoff D statistic

The preference for the NB may be biased due to the fact that different distributions have different numbers of parameters (e.g., 1, 2, 3 for the Poisson, NB and ZINB respectively). The *D* statistic provides an alternative approach that is insensitive to model complexity. Here we used a null hypothesis *H*_0_ stating that there is no difference between the cumulative distribution of the observed expression of the gene and the cumulative distribution of the theoretical distribution. For each dataset and each gene (trimmed mean of 1%), we rejected the null hypothesis using an (unadjusted) p-value of 0.01. In 12 out of the 13 datasets, the preference remained the NB distribution (Supplementary Figure S1A).

### NB versus exponential distribution

For the majority of transcripts, the NB and exponential distributions had the first and second lowest AIC scores with averages of 60.4% and 35.8% respectively (Figure 1A). We asked if the preference for NB was an artifact of how expression was pre-processed. More specifically, we note that the shape of the NB distribution when there is high dispersion *θ* and a low expected count *µ* is similar to the shape of the exponential distributions especially when there are extreme transcript counts in the right tail. Therefore, we hypothesize that relatively strong preference for the exponential distribution is due to the presence of outliers. In particular, we asked whether an increase in the trimming of outliers (especially those in the right tail) would decrease the preference for the exponential distribution. This is confirmed in Supplementary Figure S1B, further strengthening our belief that the fRNA-seq transcript count is distributed according to the NB distribution.

**Figure S1:**
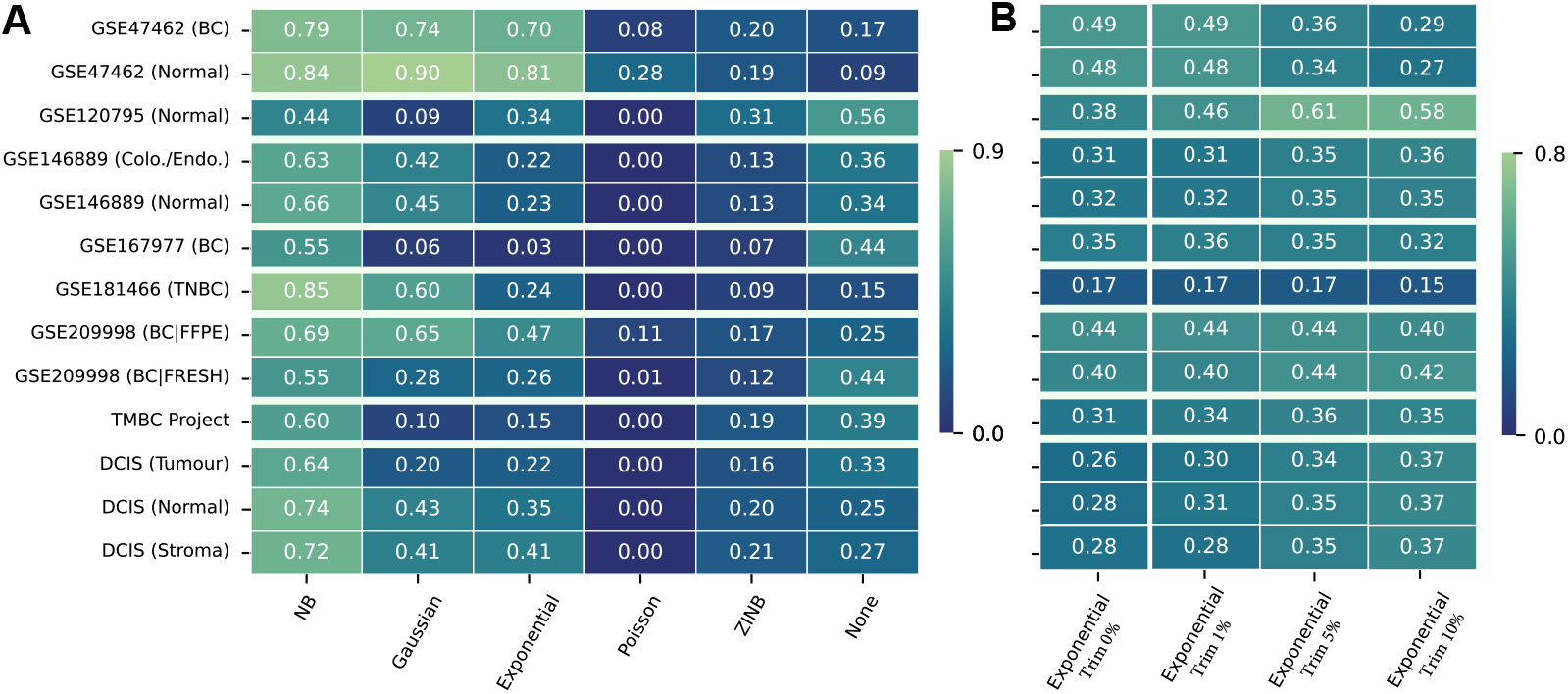
**A**. The fraction of transcripts where the null hypothesis is accepted according to the KS two-samples test. Column labelled None indicates the fraction of transcripts where the null hypothesis was not accepted for any of the distributions. **B**. The fraction of transcripts that preferred an exponential distribution (via the AIC measure) across different thresholds for mean trimming. Increased trimming reduces the number of outliers; the removal of right tail outliers reduces the preference for the exponential distribution.

**Figure S2:**
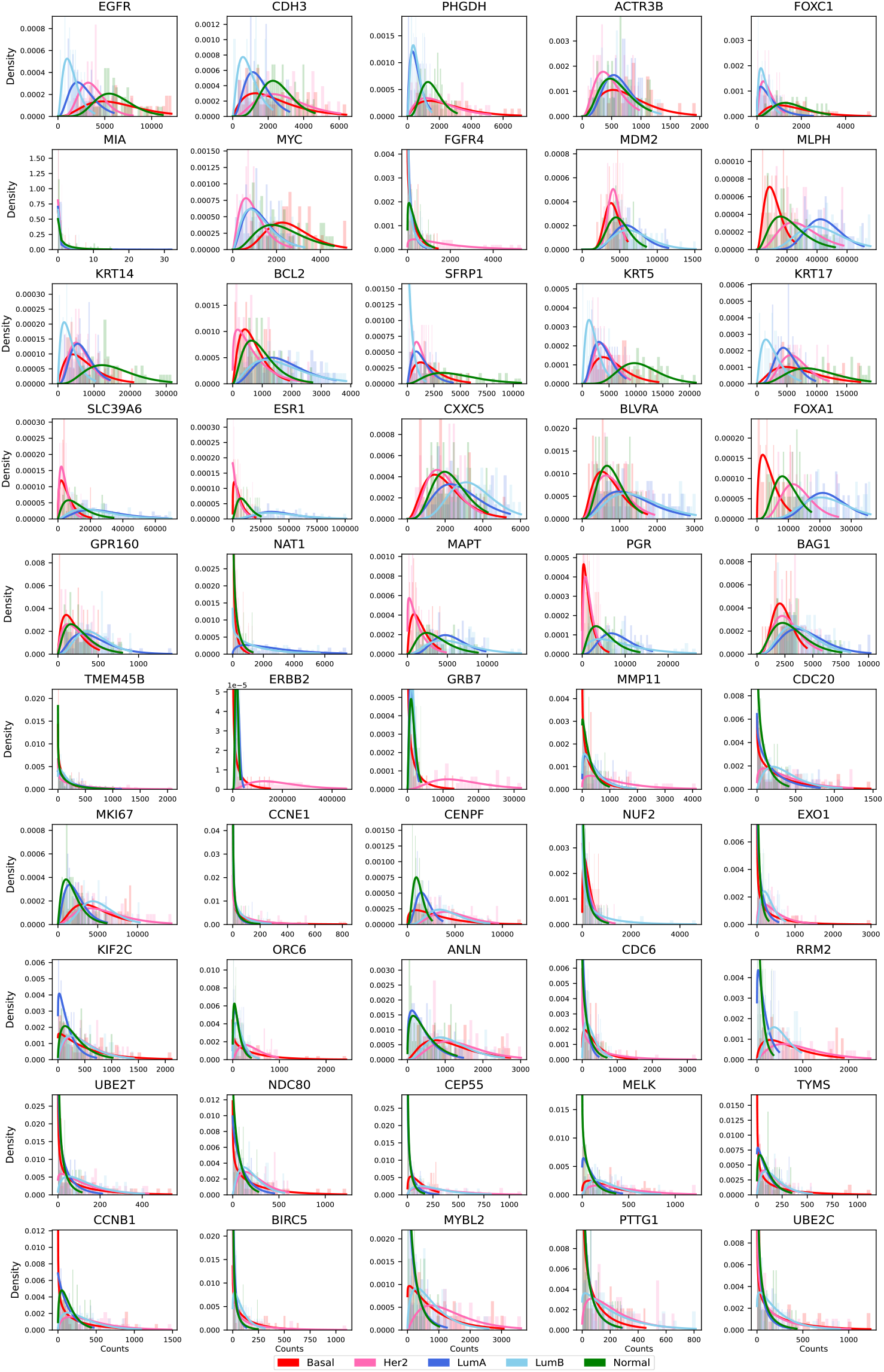
Histograms of counts for each PAM50 transcript in the Sunnybrook DCIS tumor cohort. Samples were first normalized by median library size, and subjected to outlier trimming. Samples were stratified by the PAM50 classifier before an NB distribution was fit to each transcript. The *y*-axis was truncated to improve readability.

## 4 Supplementary Methods

**Figure S3:**
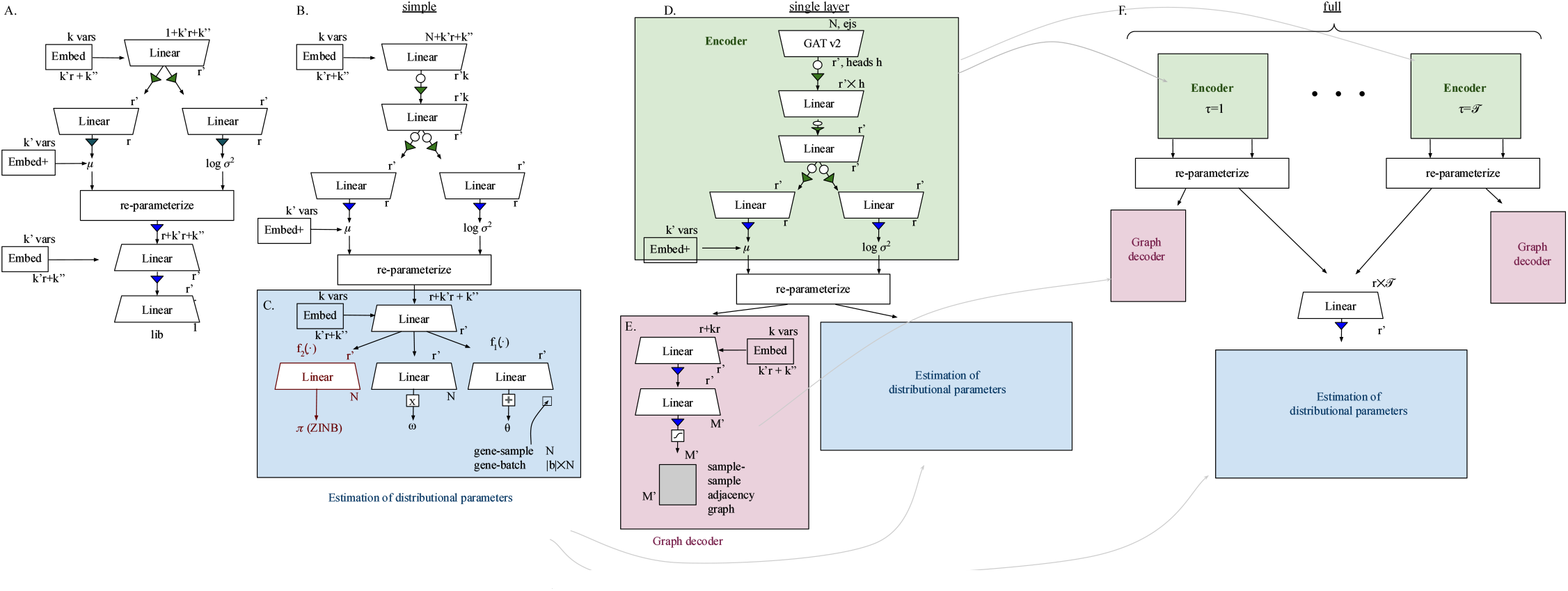
A schematic describing the neural networks. **A**. This VAE encodes sample library size. Here *k*′ and *k*″ are the number of categorical and continuous correction variables respectively, and *r* is the dimension of the latent space. The Embed symbol denotes the concatenation of *k*″ embeddings for the categorical variables with total dimension *k′′ · r*. It is concatenated to the continuous variables and the log-sample read counts. The Embed+ symbol denotes the matrix sum of the *k*″ embeddings and the latent space, which also has dimension *r*. **B**. The simple network with a single tissue and without an adjacency network. The empty circle indicates a dropout node, a green triangle indicates an ELU function and a blue triangle indicates a leaky ReLU function. *b* represents the width (in a one-hot encoding) of the adjustment variables. **C**. The decoder (blue box) generates estimates of the parameters for the negative binomial distribution with location parameter *ω* and dispersion *θ*. If ZINB is used, it also produces *π*, the probability that a variate is 0 and not generated by the *NB* distribution. The *X* and *−* indicate a *softmax* and *softplus* function, respectively. **D**. The single layer model starts with a GAT version 2 layer with *h* heads. **E**. (pink box) The decoder consists of a graph decoder used to estimate edges of the sample-sample adjacency network, and the estimators of the distribution parameters for the NB or ZINB. The output is transformed with a sigmoid function. **F**. The full model accepts *𝒯* tissues described by both count matrices and adjacency networks.

### 4.1 The PREFFECT model

We describe a set of non-linear mappings to compute variational posteriors given the expression matrix and variables (e.g., batch) for adjustment. *q*_Φ_ is the resultant estimation of the true posterior distribution *p*_Θ_, where Φ, Θ denote all relevant underlying parameters. The exact parameterization of *q*_Φ_ depends on which of the tree models (simple, single-tissue and full) is used (Supplementary Figure S3). *q*_Φ_ is composed of several component conditional VAEs described below.

The input to all models is partitioned into minibatches with row dimension *M*′ where size *M*′ ≤*M* (Supplementary Figure S3, top). Count matrices are restricted to only the *M* samples of the minibatch with width equal to the number of transcripts *N*. Optionally, (biological and technical) adjustment variables that describe the samples can also be passed. Such variables can either be categorical (with total width of their encoding denoted by *k* or continuous (with total width denoted by *k* ″). We use *K* to denote all such adjustment variables. Adjacency networks for the single tissue and full models are also partitioned into *M′ × M′* adjacency matrices. The relevant components of VAEs to handle adjacencies can also include adjustment variables *K*.

Library sizes are computed and fit using a log-Normal distribution. Across all models, the observed sample library sizes *L* can be allowed to vary during model fitting. Supplementary Figure S3A depicts the neural network underlying the encoder 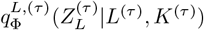 that approximates the true posterior distribution *p*_Θ_(*Z*^(*τ*)^|*L*^(*τ*)^, *K*^(*τ*)^). The encoder for the simple model is a three-layer neural network that outputs a latent space with final dimension *r* (Supplementary Figure S3B). The encoder *q*_Φ_(*Z*|*X, K*) is an estimate of the true posterior *p*_*θ*_(*Z*|*x, K*) where *θ* is the set of all parameters of the neural network. When multiple tissues are present (full model), we write 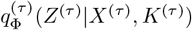.

The single tissue layer (*𝒯* = 1) and full models (*𝒯 >* 1) encode the input sample-sample adjacency networks into separate Gaussian VAEs. The encoder *q*_*A*,Φ_(*Z*_*A*_| *X, A, K*) for the networks and associated adjustment variables estimate the true posterior distribution *p*_*A,θ*_. The underlying neural network for each tissue *τ* begins with a GAT version 2 convolutional layer [52] with a tune-able parameter *h* denoting the number of heads in the attention mechanism (Supplementary Figure S3C). For each expression matrix *X*, each edge *e*_*ij*_ in the adjacency matrix *S* associated with *X*, and for each attention head 1 *≤ η ≤ h*. we compute:

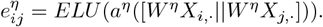

Here || denotes the concatenation operation, *W*^*η*^ is the weight matrix, and *a*^*η*^ is a vector with dimension R^1*×*2*r*^. These values are then normalized across all neighbors 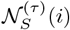 using a softmax function:

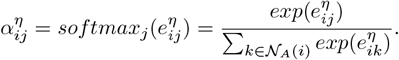

These normalized attention coefficients are then used to compute a linear combination of their features, and the *h* heads are concatenated as follows:

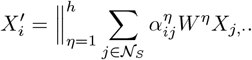

The width of the embedding is therefore *rh*.

The reparameterization step generates two sets of variates for the latent space of each tissue:

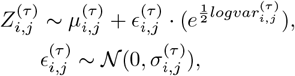

across samples 1 *≤ i ≤ M′* and for each dimension of the latent space 1 *≤ j ≤ r*. A separate decoder produces an *M ′ × M′* estimation of *Â*^(*τ*)^ for each adjacency tissue (only single-tissue and full models).

The decoder in the simple model consists of two linear layers that produce estimates of the parameters for the NB distribution including *c*_*m*_, which is a probability density function where *c*_*m,g*_ is the fraction of counts for the transcript *g* in each a sample *m*, and the inverse dispersion *θ. θ* can be estimated for each gene-sample combination (denoted *θ*_*m,g*_, or for each gene within each batch (*θ*_*g*_). Let *l*_*m*_ be the library size for sample *m* (optionally, this may be optimized during the learning step). The NB distribution is expressed as a hierarchical Gamma-Poisson model:

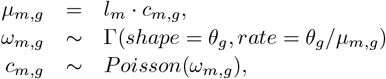

for each sample *m* and transcript *g*. See also Figure 3C. If the ZINB is preferred, the dropout probability *π* is also estimated. The final reconstructed expression is defined as follows:

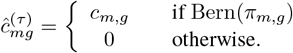

The other models have a set of latent spaces *Z*_1_, …, *Z* _*𝒯*_ which are concatenated during the decoding process (Supplementary Figure S3E):

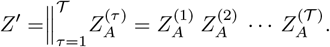

This extends the neighborhood expression learned via the attention mechanisms from a single tissue to the neighborhoods from all tissues (Figure 7).

### 4.2 Loss

We follow standard practice [57] for computing loss during training via the evidence lower bound, which combines the expected log-likelihoods of the reconstruction terms and the reverse KL divergence to regularize the posterior distributions of the latent variables. We assume that the priors follow a multidimensional normal with the identity covariance matrix. That is, they follow a *N* (0, *I*_*r*_) distribution where *I*_*r*_ is the identity matrix over the *r* dimensions of the latent space.

For the full model, this is formulated as:

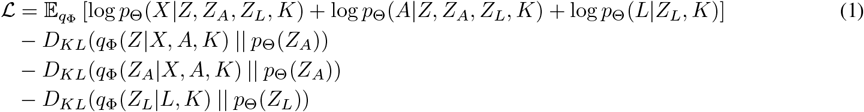

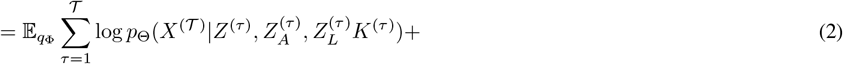

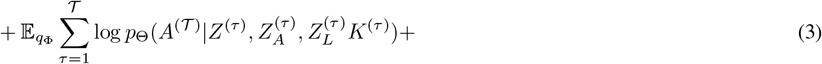

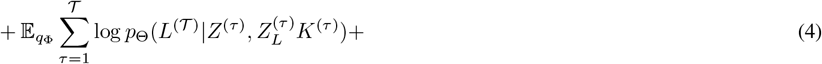

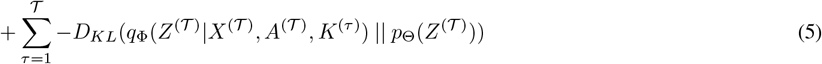

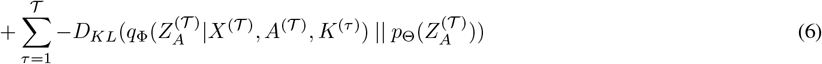

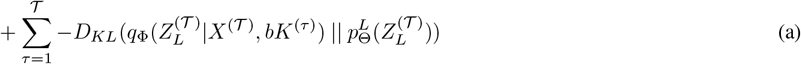

Equation 1 is computed as either *L*_*ZINB*_ = *−log*(*ZINB*(*ĉ* |*π, ω, θ*)), where *ZINB*(*X* |*π, ω, θ*) = *πδ*_0_ + (1 *−π*)*NB*(*X* |*ω, θ*) and *δ*_0_ is the Dirac function, or *L*_*NB*_ = *−log*(*NB*(*c* |*ω, θ*)), where NB is the negative binomial function. Equations 2 and 3 are computed using binary cross entropy. *D*_*KL*_ refers to the KL divergence. Equation (a) is the KL divergence if the library sizes are allowed to vary. There are also options within the PREFFECT code base to bias the learner to a disentangled latent space with respect to the adjustment variables *K* following the approach from Chen et al. [73].

